# Single-cell and spatial transcriptomics reveal a cyst-neuron-microglia network underlying brain damage during *Toxoplasma gondii* infection

**DOI:** 10.64898/2026.11.18.767391

**Authors:** Chi Zhang, Yingying Lu, Weiling Wu, Zhihao Zhou, Jiating Chen, Wenjun Huang, Min Chen, Weihao Zou, Lijuan Zhou, Shuyu Zheng, Xiangyi He, Xinyue Liao, Hongjuan Peng

## Abstract

*Toxoplasma gondii* infects approximately one-third of the world’s population, and persistent brain infection links to neuropsychiatric disorders. How cysts interact with host brain cells and tissue to sustain infection and cause neuronal damage remains unclear. We combined single-nucleus RNA sequencing with Stereo-seq spatial transcriptomics to map mouse brains across acute and chronic *T. gondii* infection. We found neurons in the cyst neighbourhood recruit interferon-responsive microglia via the APP-CD74 axis; these microglia breached the cyst wall with invasive pseudopodia, the released parasites formed secondary cysts; and microglial depletion with PLX5622 during chronic infection significantly reduced brain cyst burden, supporting that continual cyst turnover facilitates persistent infection. In mice with chronic infection, suppressed NMDA receptor signalling, enhanced complement-mediated synaptic pruning, impaired adult neurogenesis, and induced lipid reprogramming was found in the cyst neighbourhood. This cyst-neuron-microglia network thus underpins both infection persistence and neuronal injury, offering new mechanistic insights into pathogen-host interactions.

## Introduction

*Toxoplasma gondii* (*T. gondii*) is an obligate intracellular protozoan that infects humans and most warm-blooded animals, with approximately one-third of the global population infected^1,2^. Whereas infection is usually asymptomatic in immunocompetent individuals, immunocompromised patients may develop severe toxoplasmosis, including toxoplasmic encephalitis (TE) and retinochoroiditis^1,3^. After oral ingestion of oocysts or tissue cysts, released sporozoites and bradyzoites convert to tachyzoites, which disseminate through the blood and lymphatic circulation, and a proportion cross the blood–brain barrier (BBB) to enter the brain parenchyma^4,5^, where immune pressure drives their conversion into bradyzoites that establish persistent latent infection within tissue cysts^6,7^. Chronic toxoplasmosis has been associated with increased risk of neuropsychiatric disorders^8^, including schizophrenia^9,10^, Alzheimer’s disease^11,12^, Parkinson’s disease^13^, depression^14^, anxiety^15^ and cognitive decline^16^. In immunosuppressed populations, such as patients with AIDS, cancer patients undergoing chemo- or radiotherapy, and organ transplant recipients, primary or reactivated infection can cause severe organ damage and even life-threatening TE^3,17–20^; occasional cerebral toxoplasmosis accompanied by neuropsychiatric symptoms has also been reported in immunocompetent individuals^21,22^.

Traditionally, cerebral toxoplasmosis has been equated with acute fulminant TE in severely immunocompromised individuals: a necrotising encephalitis resulting from cyst reactivation and uncontrolled tachyzoite proliferation, highly lethal if untreated^18^. Accumulating evidence, however, supports a broader disease spectrum, from asymptomatic carriage to subclinical neuropsychiatric impairment and even clinical encephalitis in susceptible individuals^23,24^. Current drugs poorly cross the BBB and barely penetrate the cyst wall, leaving them largely ineffective against intracystic bradyzoites^25^; and with no indication for prophylactic treatment of chronic carriers, intervention is reserved for symptomatic patients. These clinical dilemmas converge on a long-unresolved question: how does *T. gondii* sustain persistent brain infection for decades while simultaneously inflicting neuronal damage?

The classical model holds that cysts reside primarily within neurons^4,26,27^ in a state of metabolic quiescence and immunological inertness, with neuropathology arising mainly from acute-phase inflammation. Chronic infection is, however, essentially dynamic: bradyzoites maintain low-level replication within cysts^28^; ruptured cysts account for approximately 0.27% of all cysts and are accompanied by focal microglial and monocyte/macrophage infiltration^29,30^; and *in vivo* two-photon imaging has captured CD11c^+^ antigen-presenting cells (APCs) near cysts beginning to rupture^31^. Chronic infection thus reflects a dynamic equilibrium between parasite proliferation and immune control^32^, with the number of infected neurons still increasing in the early chronic phase^27,33^. Moreover, behavioural and neuropathological abnormalities correlate with brain cyst burden and inflammation severity^30^. The central question thus becomes what drives cyst rupture and re-formation, and how this cycle progressively damages neurons.

Two pathways of cyst rupture are proposed. In the immune-mediated pathway, although cytotoxic T lymphocyte-mediated lysis of infected cells does not kill intracellular *T. gondii*^34^, CD8^+^ T cells reduce brain cyst numbers through perforin secretion and confer protection during chronic infection^35^. Monocytes, macrophages and microglia accumulate at rupture sites^29,36^, and human (but not rodent) cytolytic lymphocytes express granulysin, which acts with perforin and granzymes to kill intracellular parasites^37^. In the endogenous pathway, cysts in dopaminergic neurons can alter host dopamine metabolism via parasite-derived tyrosine hydroxylase^38,39^, and cyst growth may disrupt host homeostasis, causing endoplasmic reticulum (ER) stress or mitochondria-dependent membrane lysis^40^ and thereby triggering egress. The relative contributions of these two pathways *in vivo* remain unclear.

Because murine cytotoxic T cells destroy cyst-harboring host cells but fail to kill the enclosed parasites^34,35^, clearance of exposed cysts depends on other mechanisms. Mononuclear phagocytes and microglia cluster at rupture sites and may themselves be responsible for cyst rupture^29,36^. *In vitro*, released bradyzoites are highly motile, can escape host cells without destroying them, and can even form new cysts directly, skipping the tachyzoite stage^28^; whether free bradyzoites re-infect neural cells and spread to distant parenchyma is largely unknown. On the host side, infected neurons present antigen via major histocompatibility complex (MHC) class I to keep infection in check^41^, and brain-resident CD8^+^ T cells act against *T. gondii*^42^; whether cytotoxic T cells sense neurons harboring mature cysts remains untested, although rupture of cysts surrounded by APCs has been observed^31^. *T. gondii* can also inject effector proteins into never-invaded cells or deliver parasite-derived molecules via extracellular vesicles^43,44^.

How chronic infection correlates with neuropsychiatric disease remains far from fully understood^45^. Cysts reside mainly in neurons rather than glial cells^27,46^, and bradyzoites can drive dopamine release from dopaminergic neurons, producing marked behavioural changes^38,39^; *in vitro*, rising parasite burden reduces the dendritic complexity of primary neurons^44^. Beyond this, infection compromises the BBB and the choroid plexus barrier^47–50^ with focal inflammation and neurovascular unit dysfunction^51^; reduces neuronal fibre density, dendritic complexity, spine density and synaptic proteins^52^ and impairs fear memory consolidation^53^; and causes tau hyperphosphorylation with hippocampal neuron apoptosis^54^, elevated Aβ with reduced N-methyl-D-aspartate receptor (NMDAR) expression^55^, and IFN-γ-driven IDO activation with tryptophan depletion and reduced serotonin synthesis, triggering depressive-like behaviour^56,57^. Immune-side mechanisms include Caspase-8-dependent restraint of brain parasite burden by CD8^+^ T cells^58^, IL-1-driven hippocampal DNA double-strand breaks and cognitive deficits^59^, microglial and mononuclear-phagocyte IFN-γ production^60^, microglial participation in neuronal apoptosis during TE^61^, and downregulation of astrocyte GLT-1 by vesicles from cyst-harboring neurons, causing excitotoxicity^44,62^. However, how cysts, neurons and immune cells interact *in situ* to drive these processes remains poorly resolved at single-cell and spatial resolution.

In this study, we combined single-nucleus RNA sequencing (snRNA-seq) with high-resolution Stereo-seq spatial transcriptomics to characterize, *in situ*, the transcriptional landscape and immune microenvironment of the infected mouse brain, from the whole-brain level to the cyst neighbourhood. By integrating these data with cellular and animal pathology experiments, we found that *T. gondii* sustains persistent central nervous system (CNS) infection through a “cyst–neuron–microglia interaction network”; the accompanying neuronal injury manifests as dampened synaptic activity and plasticity in mature neurons and as compromised adult neurogenesis, with a consequent depletion of immature neurons.

## Results

### A single-nucleus and spatially integrated atlas of the *T. gondii*-infected mouse brain

To resolve the CNS response to *T. gondii* infection at single-cell and spatial resolution, we combined snRNA-seq with Stereo-seq spatial transcriptomics (Figs. 1 and 2), sampling uninfected mice (0 days post-infection (dpi)) and both the acute (12 dpi) and chronic (60 dpi) phases of infection.

**Fig. 1.**
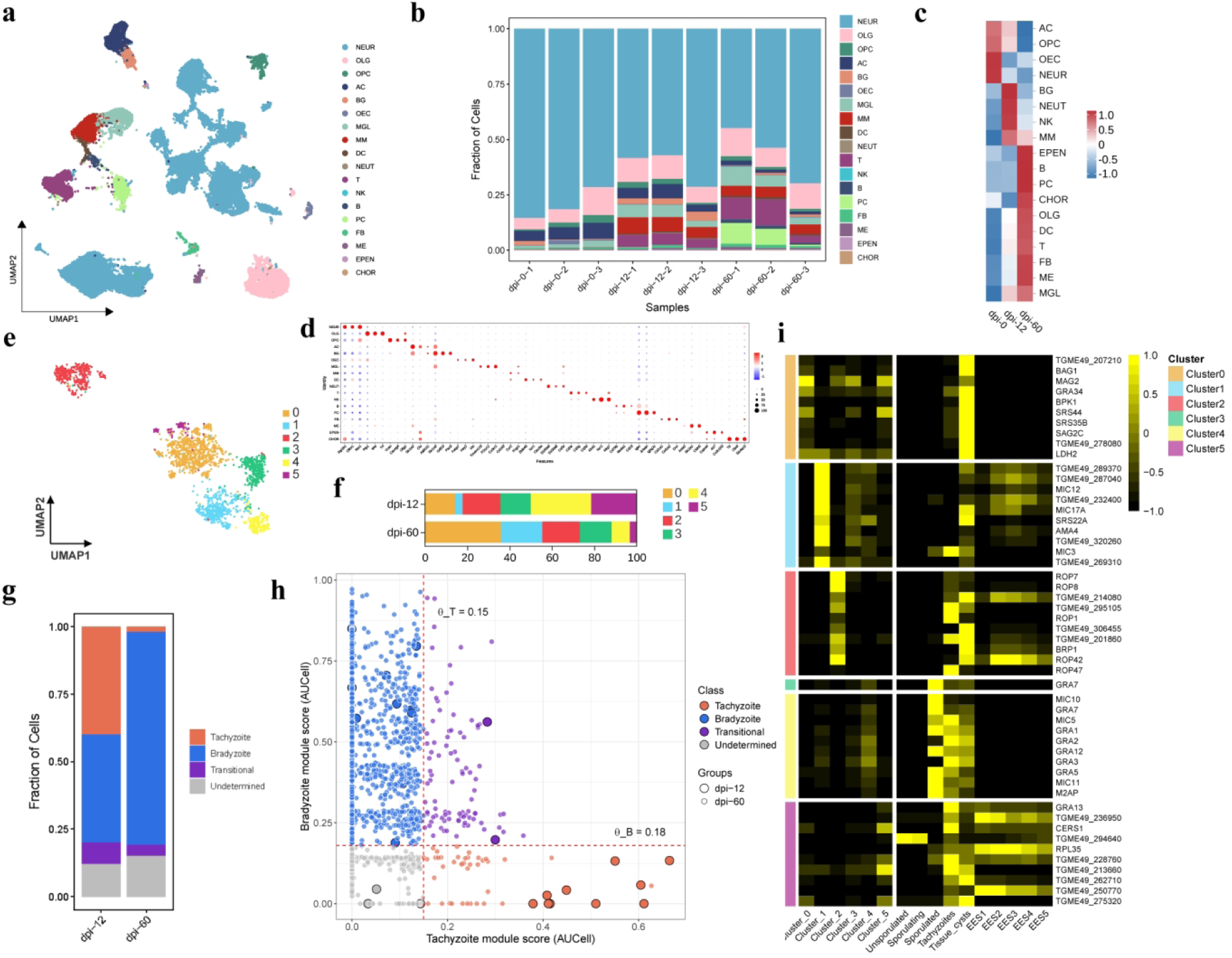
Single-nucleus transcriptomic atlas of the *T. gondii*-infected mouse brain and intracerebral parasite heterogeneity. a, UMAP of 79,735 host nuclei at 0, 12 and 60 dpi, coloured by 18 cell populations. NEUR, neurons; OLG, oligodendrocytes; OPC, oligodendrocyte precursor cells; AC, astrocytes; BG, Bergmann glia; OEC, olfactory ensheathing cells; MGL, microglia; MM, monocytes/macrophages; DC, dendritic cells; NEUT, neutrophils; T, T cells; NK, natural killer cells; B, B cells; PC, plasma cells; FB, fibroblasts; ME, mural/endothelial cells; EPEN, ependymal cells; CHOR, choroid plexus epithelial cells. b, Cell-population fractions per sample (n = 3 mice per time point). c, Row-scaled cell-population frequencies by infection stage. d, Canonical markers used for cell-population annotation (dot size, percentage of expressing cells; colour, scaled mean expression). e, UMAP of 2,188 parasite nuclei (12 dpi, n = 28; 60 dpi, n = 2,160), coloured by six subclusters. f, Parasite subcluster composition at 12 and 60 dpi. g, Parasite stage composition. Tachyzoite (T) and bradyzoite (B) module AUCell scores were calculated for 2,188 parasites; 42 cells without detected module genes were excluded, leaving 2,146. Cells were classified using θ_T_ = 0.15 and θ_B_ = 0.18 as Tachyzoite, Bradyzoite, Transitional (T^+^B^+^) or Undetermined (Methods; Supplementary Tables 1 and 5; Supplementary Fig. 1m–r). Bars show composition at 12 dpi (n = 25 classified cells) and 60 dpi (n = 2,121). h, T versus B module scores for the 2,146 classified parasites. Circled symbols, 12-dpi cells; dashed lines, thresholds. Module genes, classification rules and per-cell scores are provided in Methods and Supplementary Tables 1 and 5. i, Top 10 differentially expressed genes per parasite subcluster, with reference life-cycle stages from ToxoDB (unsporulated, sporulating and sporulated oocysts, tachyzoites, tissue cysts and enteroepithelial stages EES1–5), showing scaled mean expression. Subcluster 3 has one borderline DEG, GRA7 (FDR = 0.05297), which does not pass FDR < 0.05.

**Fig. 2.**
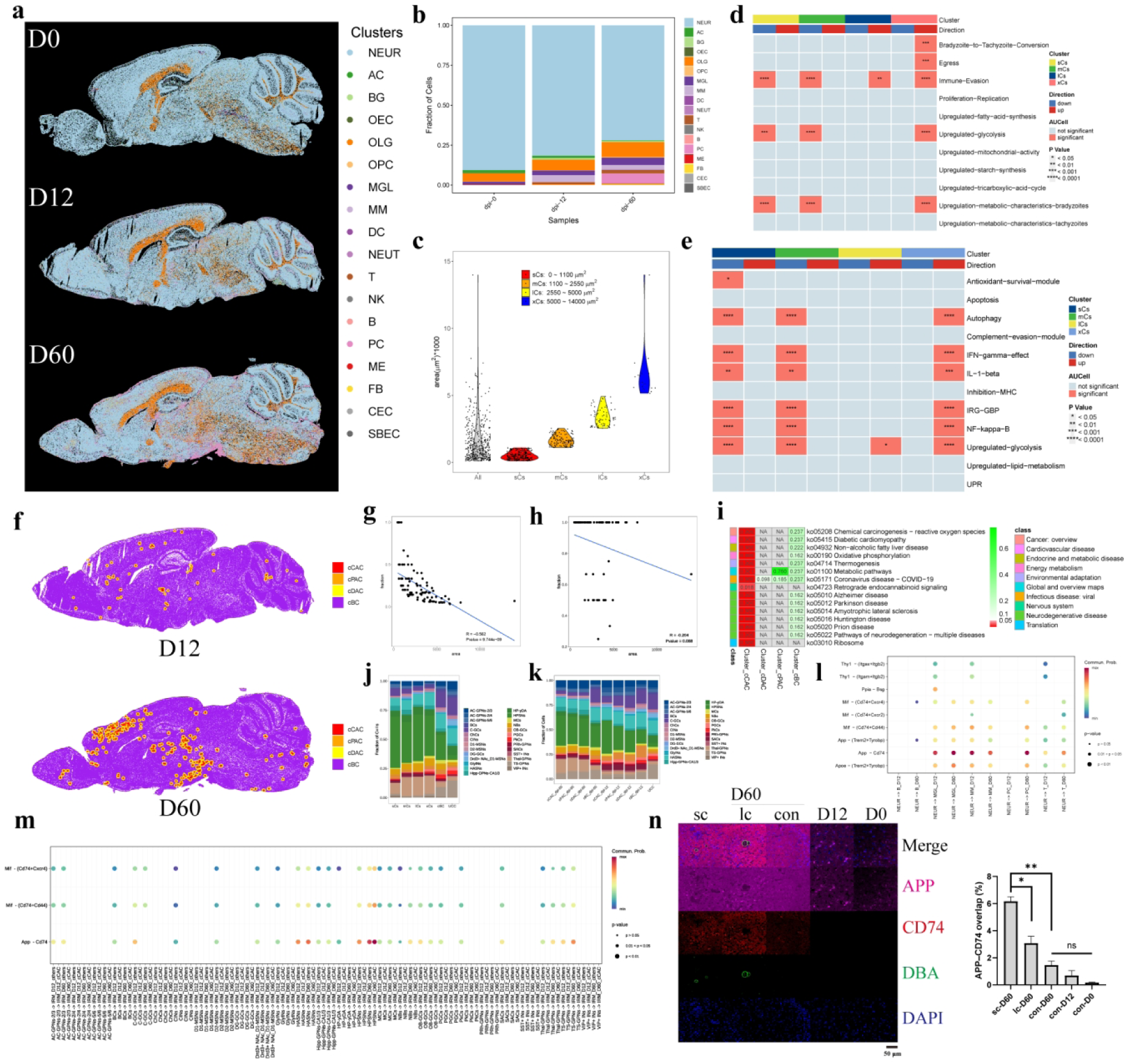
Spatial organisation of PV/cyst structures and the immune microenvironment of the cyst neighbourhood. a, Spatial deconvolution maps of 18 host cell populations on sagittal brain sections at 0, 12 and 60 dpi. b, Host cell-population fractions in the Stereo-seq data by stage. c, Cross-sectional areas and K-means size classification of PV/cyst structures (k = 4): sCs, 0–1,100 μm²; mCs, 1,100–2,550 μm²; lCs, 2,550–5,000 μm²; xCs, 5,000– 14,000 μm² (140 structures at 12 dpi; 300 at 60 dpi). d,e, Parasite gene-module AUCell scores by size class: developmental/metabolic modules (d) and host-interaction/immune-evasion modules (e). Pink, significant size association; Direction, upregulation (red) or downregulation (blue) with increasing size; *P < 0.05, **P < 0.01, ***P < 0.001, ****P < 0.0001. f, CN layers at 12 and 60 dpi: core-associated (cCAC), proximal (cPAC), distal (cDAC) and structure-non-associated background cells (cBC). g,h, Microglial fraction in the CN (g) and interferon-responsive microglial fraction among CN microglia (h) versus structure area (Pearson correlation; g, R = −0.562, P = 9.744 × 10^-^⁹; h, R = −0.204, P = 0.088). i, KEGG enrichment of IRM DEGs across cCAC, cPAC, cDAC and cBC. j, Neuronal subcluster composition by structure size class. k, Neuronal subcluster composition across CN layers at 12 and 60 dpi, with cBC and UCC as references. l, CellChat ligand–receptor analysis of neuron-to-immune-cell communication (colour, communication probability; dot size, P-value category: P ≥ 0.05, 0.01 < P < 0.05, P < 0.01). m, CellChat analysis of neuronal-subcluster-to-IRM communication in cCAC versus other layers at 12 and 60 dpi. n, APP, CD74, DBA and DAPI immunofluorescence around small cysts (sc, < 40 μm diameter), large cysts (lc) and matched cyst-free controls (con) at 60 dpi, and controls at 12 and 0 dpi. Right, APP–CD74 overlap (n = 3 mice per group; *P < 0.05, **P < 0.01; ns, not significant).

**Table 1.** Mobile-phase gradient.

| **Time (min)** | **Flow rate (mL/min)** | **Phase A (%)** | **Phase B (%)** | **Curve** |
|---|---|---|---|---|
| Initial | 0.2 | 60 | 40 | 6 |
| 4.00 | 0.2 | 57 | 43 | 6 |
| 4.10 | 0.2 | 50 | 50 | 6 |
| 24.00 | 0.2 | 46 | 54 | 6 |
| 24.10 | 0.2 | 30 | 70 | 6 |
| 36.00 | 0.2 | 1 | 99 | 6 |
| 40.00 | 0.2 | 1 | 99 | 6 |
| 40.10 | 0.2 | 60 | 40 | 6 |
| 44.00 | 0.2 | 60 | 40 | 6 |

**Table 2.** Mass-spectrometry parameters.

| **Parameter** | **Value** |
|---|---|
| Capillary voltage | 2.30 kV |
| Cone voltage | 40 V |
| Source offset | 10 V |
| Source temperature | 150 °C |
| Desolvation temperature | 500 °C |
| Cone gas flow | 50 L/h |
| Desolvation gas | 6.0 bar |
| Reference capillary voltage | 2.20 kV |
| Acquisition mode | MSE |
| Low collision energy | 6 eV |
| High collision energy | 30–50 eV |
| Lock mass | Leucine enkephalin, 0.2 µg/mL |
| Mass-axis calibration | Sodium formate, 0.5 mmol/L |

Histological examination confirmed parasite invasion and inflammatory infiltration at these time points (Supplementary Fig. 1a). After quality control, we obtained 79,735 host nuclei and 2,188 parasite nuclei and identified 18 host cell clusters (Fig. 1a): neurons (NEUR), oligodendrocytes (OLG), oligodendrocyte precursor cells (OPC), astrocytes (AC), Bergmann glial cells (BG), olfactory ensheathing cells (OEC), microglia (MGL), monocytes/macrophages (MM), dendritic cells (DC), neutrophils (NEUT), T lymphocytes (T), natural killer cells (NK), B lymphocytes (B), plasma cells (PC), fibroblasts (FB), mural/endothelial cells (ME), ependymal cells (EPEN) and choroid plexus epithelial cells (CHOR) (markers in Fig. 1d). Further subclustering yielded 80 subclusters in total (Supplementary Note 1, Supplementary Fig. 1b–k).

Cell frequencies changed markedly across infection stages (Fig. 1b,c; statistical tests in Supplementary Fig. 1l). Neurons and astrocytes decreased significantly over the course of infection, whereas oligodendrocytes increased; NK cells and neutrophils increased sharply in the acute phase; B cells and plasma cells rose only in the chronic phase; T cells and DCs increased progressively; microglia and monocytes/macrophages peaked during the acute phase and then declined gradually; and OPCs declined significantly in the chronic phase (all *P* < 0.0001).

We also profiled the parasites themselves and identified six parasite subclusters (fewer parasite nuclei were captured at the acute phase; Fig. 1e). Their proportions shifted across stages (Fig. 1f): subclusters 0 and 1 became dominant in the chronic phase, whereas subclusters 4 and 5 declined. Tachyzoites, bradyzoites and transitional-stage parasites coexisted at 12 dpi; by 60 dpi, bradyzoites predominated (Fig. 1g,h; Supplementary Fig. 1m–r; Supplementary Table 1).

Pseudotime analysis revealed a continuous differentiation trajectory across subclusters: subcluster 1 occupied the start of the axis with differentiation plasticity, and 12-dpi parasites had already acquired a chronic-stage-like pattern (Supplementary Fig. 1s–v). Subcluster 0 represented classical quiescent bradyzoites (BAG1, MAG2, GRA34, BPK1, SRS44), characterised by cyst wall and matrix components together with downregulated metabolism and stress-resistance and immune-evasion programs. Subclusters 1 (MIC12, MIC17A, SRS22A, AMA4, MIC3) and 4 (MIC10, GRA7, MIC5, GRA1, GRA2) expressed distinct microneme and dense-granule programs, the latter maintaining the cyst wall and intracystic tubulovesicular network. Subcluster 2 combined cyst-wall maintenance with a rhoptry program (ROP7, ROP8, ROP1, BRP1, ROP42); subcluster 3 was GRA7^+^; and subcluster 5 (GRA13, CERS1, RPL35, SND1, RPL26) appeared to be slowly proliferating bradyzoites (Fig. 1i).

### Cyst-neighbourhood neurons recruit interferon-responsive microglia via APP– CD74

Deconvolving the Stereo-seq data against our snRNA-seq reference mapped the 18 host cell clusters onto brain sagittal sections at each stage (Fig. 2a), with cell-frequency trajectories tracking closely between the two omics layers (Fig. 2b). We localised parasitophorous vacuoles (PVs) and tissue cysts from the on-chip ssDNA signal and identified 140 PV/cyst structures at 12 dpi and 300 at 60 dpi; these comprised PVs, cysts and mixed-stage cysts (mixCysts), together with foci that could not be confidently staged (Unresolved; 104 at each time point); among classifiable foci, PVs predominated at 12 dpi and cysts predominated at 60 dpi (Supplementary Fig. 2a,b; Supplementary Table 2). K-means clustering by cross-sectional area separated them into four size classes (labelled sCs, mCs, lCs and xCs; Fig. 2c; size ranges in Methods; spatial distribution in Supplementary Fig. 2c). The cross-sectional area of these structures correlated positively with detected transcript number (linear regression; Supplementary Fig. 2d: transcripts, R = 0.722, P = 5.84 × 10^-^⁶²; genes, R = 0.745, P = 6.47 × 10^-^⁶⁸). The acute stage was dominated by small structures, whereas all size classes occurred chronically (Supplementary Fig. 2e), indicating progressive enlargement and continued emergence of new small structures (mostly cysts) during the chronic stage, even without peripheral reinfection. Stage composition within the size classes tracked this shift (Supplementary Fig. 2f; Supplementary Table 2): PVs accounted for most classifiable structures at 12 dpi, whereas 60-dpi structures were dominated by cysts, with small chronic structures being almost exclusively cysts (92% of classifiable sCs). The mixCyst fraction rose steeply with structure size at 60 dpi, from 4% of classifiable sCs to 13%, 58% and 92% for mCs, lCs and xCs, respectively, indicating that tachyzoite-module signal accumulates as chronic cysts enlarge; such mixed-stage cysts may reflect tachyzoites co-existing with bradyzoites within the same growing cyst, or transitional-stage parasites emerging during cyst growth. Gene-set scoring revealed a negative relationship between structure size and immune activation (Fig. 2d,e): small structures were relatively poor at resisting host immunity, whereas large structures carried a stronger immune-evasion signature, i.e., structures that persist become progressively better at evading immunity.

To define the cyst microenvironment, we delineated the cyst neighbourhood (CN), defined here as the host-cell microenvironment that forms around PV/cyst structures as PVs progressively develop and remodel into cysts. Host cells were stratified by distance to the structure (CN core-, proximal- and distal-associated cells: cCAC, cPAC and cDAC; structure-non-associated background cells, cBCs; uninfected control cells, UCCs; definitions in Methods; spatial mapping in Fig. 2f). All neighbourhood and size-based analyses were performed on the full set of PV/cyst structures, which at 60 dpi were predominantly cysts. Against the cBC baseline, microglia and monocytes/macrophages were markedly enriched in the CN, most prominently early in infection (Supplementary Figs. 2g and 3d); among microglial subclusters, interferon-responsive microglia (IRMs) and pro-inflammatory microglia (PIMs) were enriched (Supplementary Fig. 2h). Microglial frequency scaled inversely with structure size (Fig. 2g), linking newly formed structures to microglial accumulation; IRMs constituted the largest microglial fraction, with a non-significant trend towards enrichment around smaller structures (Fig. 2h; R = −0.204, P = 0.088). The closer CN-associated IRMs were to a structure, the more strongly their differentially expressed genes (DEGs) were enriched for oxidative-stress and neurodegenerative-disease pathways (Fig. 2i), and the more they were associated with cytotoxic CD8^+^ T cell chemotaxis, activation and differentiation (gene set variation analysis (GSVA); Supplementary Fig. 2i), suggesting that IRMs sense the structure and recruit T cells. In the chronic brain, small-structure cores (cCACs) were dominated by IRMs, whereas large structures were ringed by more PIMs and proliferating microglia (PMs), whose DEGs were enriched in neurodegenerative-disease and cell-junction pathways (Supplementary Fig. 2j–l).

**Fig. 3.**
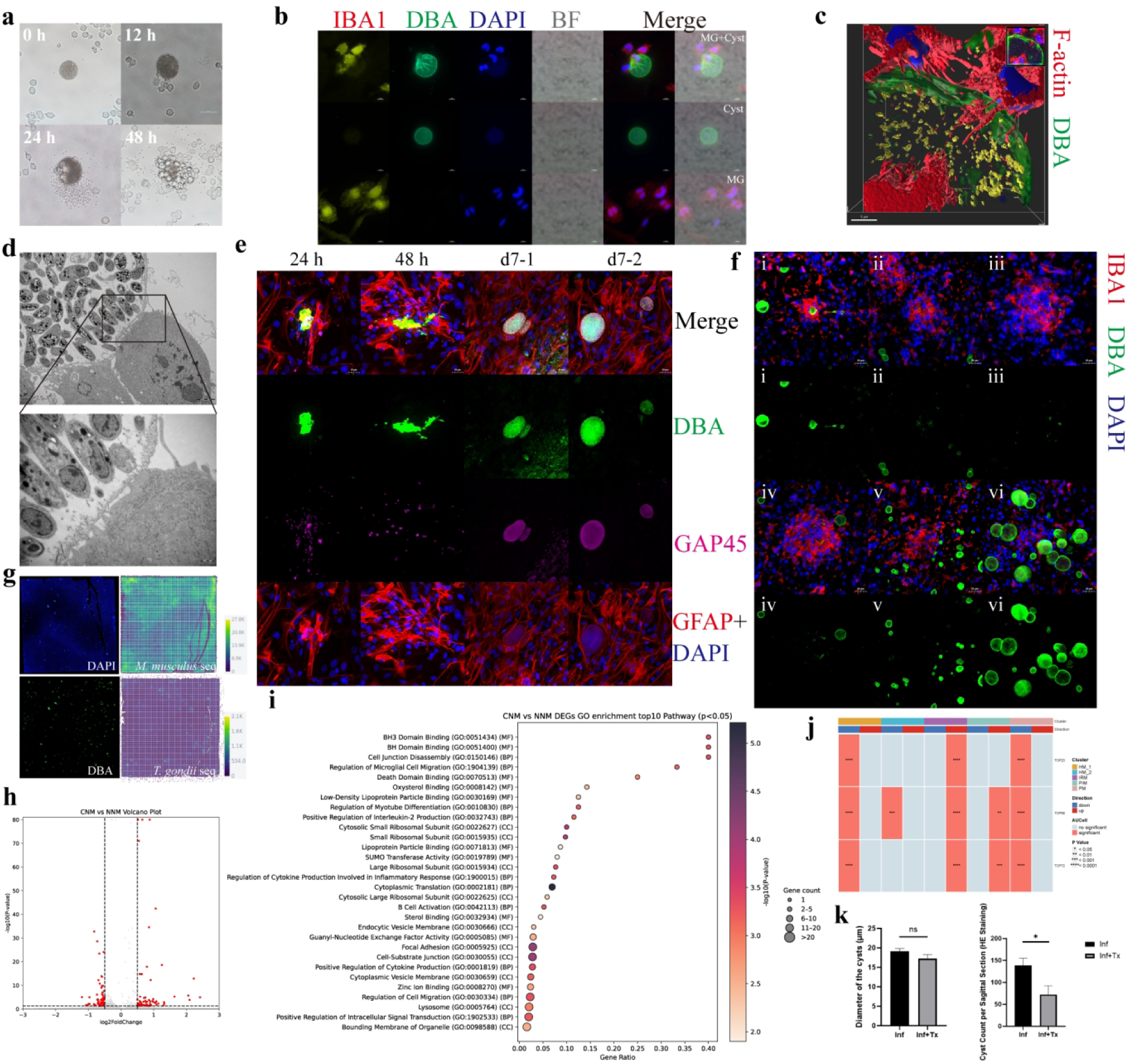
Microglia disrupt cysts and seed new cyst formation. a, Bright-field time course of BV2 microglia co-cultured with purified ME49 cysts (0, 12, 24 and 48 h). b, IBA1 (yellow), DBA (green) and DAPI (blue) immunofluorescence in primary-microglia–cyst co-cultures (MG+Cyst), with cyst-only and microglia-only controls; bright-field/merged views are included. c, N-SIM three-dimensional reconstruction of a microglial cell attacking a cyst: phalloidin/F-actin (red), DBA (green), parasite nuclei (yellow) and microglial nuclei (blue); boxed region magnified at top right. d, Transmission electron microscopy of a microglial protrusion breaching the cyst wall; the boxed region is magnified below. e, Cyst-reseeding assay in primary astrocytes at 24 h, 48 h and day 7 (two day-7 fields): phalloidin/F-actin (red), DBA (green), TgGAP45 (magenta) and DAPI (blue). f, Corresponding in vivo stages during chronic infection (i–vi): microglia approaching a cyst (i), nodules containing cyst-wall debris (ii–iv), and nascent cysts around nodules (v–vi). Merged/single-channel views show IBA1 (red), DBA (green) and DAPI (blue); representative fields of a sporadically captured process. g, Stereo-cell analysis of primary-microglia–cyst co-cultures: DAPI/DBA fluorescence and on-chip mouse/parasite transcript maps. h, DEGs between cyst-neighbourhood (CNM) and non-neighbourhood microglia (NNM) (Wilcoxon rank-sum test; |log_2_ fold change| > 0.5 and P < 0.05; 72 upregulated, 65 downregulated; Supplementary Table 3). i, Top 10 GO terms per category for CNM-versus-NNM DEGs (clusterProfiler hypergeometric test, Benjamini–Hochberg-adjusted P < 0.05). j, AUCell scores for TOP23, TOP56 and TOP72 gene sets (Supplementary Table 3) across in vivo microglial subclusters. Red, significant deviation; Direction, upregulation (red) or downregulation (blue); ***P < 0.01, ****P < 0.001, P < 0.0001. k, PLX5622 microglial depletion during months 1–2: cyst diameter (left; P = 0.1662, t = 1.576, df = 5.977) and cysts per sagittal H&E section (right; P = 0.0403, t = 2.620, df = 5.888) in infected untreated (Inf, n = 3) and infected PLX5622-treated (Inf+Tx, n = 5) mice (Welch’s unpaired t-test; mean ± s.e.m.; ns, not significant; *P < 0.05).

Highly plastic spiny neurons (HPSNs) were enriched in the CN (Fig. 2j,k) but declined markedly after infection (Supplementary Fig. 2m). To explain both HPSN loss and microglial accumulation around cysts, we performed cell–cell communication analysis between neurons and immune cells. Neurons were predicted to signal to microglia most strongly through the App–Cd74 ligand–receptor pair (Fig. 2l); during chronic infection, App–Cd74 communication between HPSNs and IRMs was strongest, particularly within the structure core (cCAC) (Fig. 2m). Immunofluorescence confirmed that APP and CD74 proteins were upregulated upon infection and increased further in the CN (*P* < 0.01), with the strongest signal around small cysts (< 40 μm in diameter; *P* < 0.05) (Fig. 2n).

### Microglia disrupt cysts and seed new cyst formation

To test whether brain-resident microglia directly disrupt cysts, we co-cultured purified cysts with a microglial cell line (BV2) (Fig. 3a) or primary microglia (Fig. 3b); in both systems microglia disrupted cysts in a contact-dependent manner. Super-resolution (N-SIM) and transmission electron microscopy (TEM) showed that microglia extended invasive protrusions that breached the cyst wall, releasing the bradyzoites within (Fig. 3c,d). To test whether this disruption drives new cyst formation, we added primary microglia and cysts to coverslips pre-seeded with confluent primary astrocytes; at 24 h, 48 h or 7 days the co-cultures were washed three times with PBS and then fixed. Released bradyzoites invaded astrocytes within 24–48 h and, by day 7, had developed into new cysts (Fig. 3e). Immunofluorescence of chronically infected brains captured the corresponding *in vivo* process (Fig. 3f): microglia recruited toward the cyst (i), cyst-wall debris within post-disruption microglial nodules (ii–iv), and new small cysts emerging around the nodules and growing (v–vi).

To define the molecular identity of cyst-disrupting microglia, we co-cultured primary microglia with cysts on Stereo-cell chips and, 6 h later, performed on-chip single-cell transcriptomic analysis of the recovered cells (Fig. 3g); nearly all recovered cells were primary microglia (Supplementary Fig. 3a). DEG analysis of manually selected cyst-associated microglia identified 72 significantly upregulated and 65 significantly downregulated genes when comparing cyst-neighbourhood microglia (CNM) with non-neighbourhood microglia (NNM) (Fig. 3h, Supplementary Table 3).

GO enrichment showed that CNM were significantly enriched for anti-apoptotic, pro-migration, pro-inflammatory and protrusion-mediated invasion pathways (Fig. 3i); for cyst-rupturing events, the altered genes were mainly enriched in environmental sensing, metabolic reprogramming and attack/degradation functions. To test whether cyst-disrupting microglia exist *in vivo*, we ranked the *in vitro* DEGs by significance into three sets (TOP23, TOP56 and TOP72) and used AUCell to search *in vivo* microglial subpopulations for a matching phenotype: the IRM subpopulation best matched the cyst-disrupting phenotype (Fig. 3j).

Finally, to verify that microglial cyst disruption is critical for cyst persistence *in vivo*, we depleted microglia from chronically infected mice with the colony-stimulating factor 1 receptor (CSF1R) inhibitor PLX5622. We first quantified cyst diameter and number over time: in our model (129S2 mice orally infected with ME49), cyst numbers rose during months 1–2 of infection and declined by month 6 (Supplementary Fig. 3b). PLX5622 was therefore given from month 1 to month 2, with sampling at 2 months. Microglia-depleted mice showed no significant difference in cyst diameter but significantly fewer cysts (Fig. 3k), corroborating a key role of microglia in cyst maintenance and disease progression in cerebral toxoplasmosis.

### Infection with *T. gondii* causes blood–brain barrier dysfunction, excessive synaptic pruning and impaired adult neurogenesis

Multiple studies have shown that *T. gondii* infection compromises the BBB in mice^47–50^. We quantified peripheral immune cell adhesion to the BBB by proximity scoring (cells with a proximity score to endothelial cells greater than 0.414 were defined as BBB-adherent; Supplementary Fig. 3c). BBB residence and parenchymal infiltration were stage-specific: mononuclear phagocytes, neutrophils and T cells adhered and infiltrated prominently early, whereas in the chronic phase DCs and B cells, particularly plasma cells, were additionally recruited (Fig. 4a). We further defined the 10% of astrocytes closest to endothelial cells as BBB-associated astrocytes (Fig. 4b): their activation gene-set score increased after infection (Fig. 4c), while their endfoot function score decreased persistently (Fig. 4d), suggesting that infection-induced astrocyte activation compromises BBB maintenance.

**Fig. 4.**
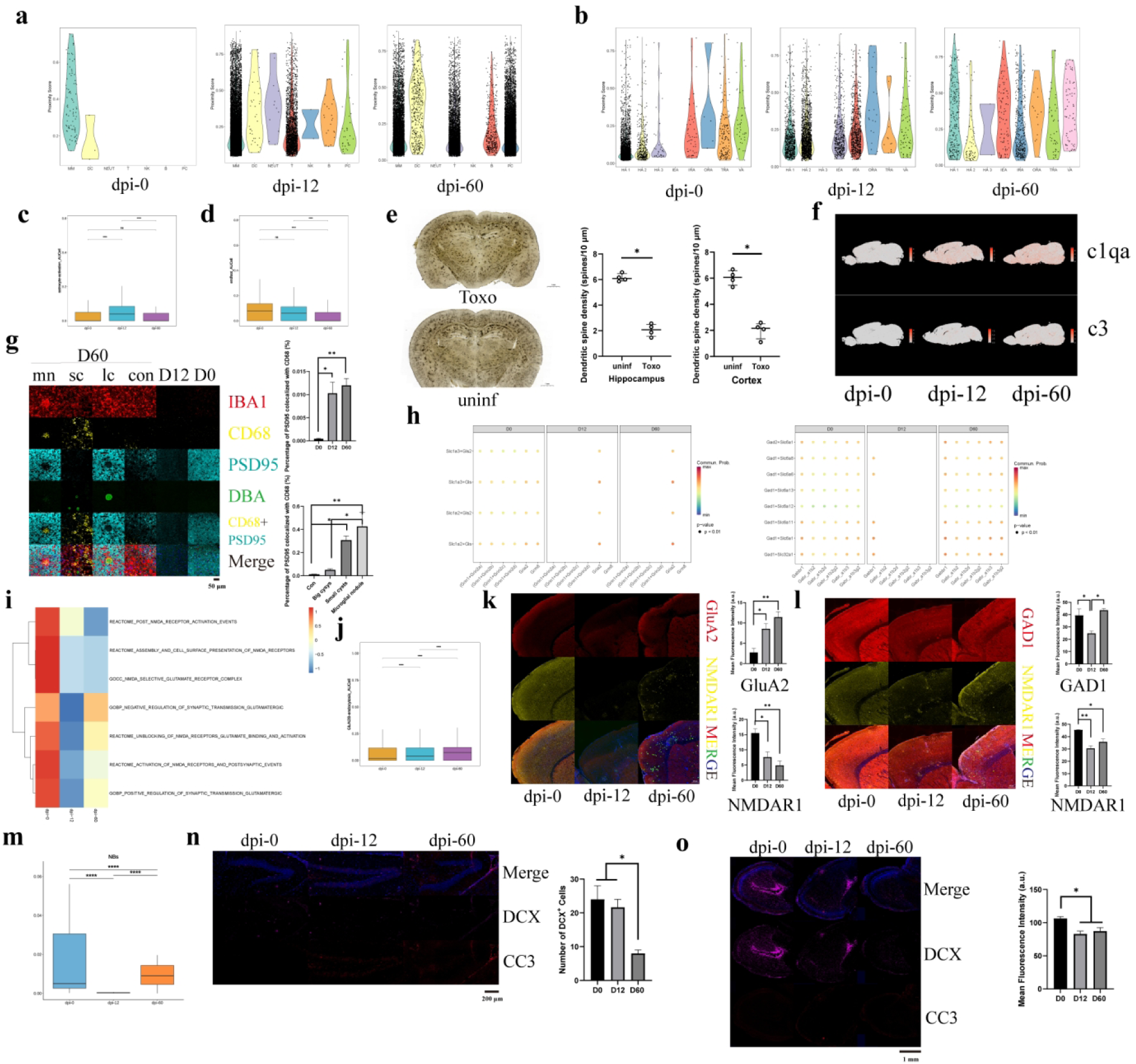
Blood–brain barrier dysfunction, excessive synaptic pruning and impaired adult neurogenesis during chronic infection. a,b, Proximity scores between endothelial cells and peripheral immune populations (a) or astrocyte subclusters (b) at 0, 12 and 60 dpi. Peripheral cells with proximity > 0.414 were defined as BBB-adherent; the 10% of astrocytes closest to endothelial cells were defined as BBB-associated. c,d, Astrocyte activation (c) and endfoot-function (d) AUCell scores in BBB-associated astrocytes. e, Golgi staining and dendritic spine density in hippocampus and cortex of uninfected and chronically infected mice (n = 4 mice per group; Mann–Whitney test, *P = 0.0286 for both regions; Welch’s unpaired t-test confirmation, P < 0.0001; mean ± s.e.m.). f, Stereo-seq feature maps of C1qa and C3 at 0, 12 and 60 dpi. g, Cortical IBA1, CD68, PSD95, DBA, CD68–PSD95-colocalisation and merged images around microglial nodules (mn), small cysts (sc, < 40 μm diameter), large cysts (lc) and cyst-free parenchyma (con), plus 12- and 0-dpi controls. Right, percentage of PSD95 colocalised with CD68 across stages (top) and 60-dpi regions (bottom; n = 3 mice per group). h, CellChat analysis of glutamatergic (left) and GABAergic (right) ligand–receptor communication among neurons (colour, communication probability; dot size, P-value category: P ≥ 0.05, 0.01 < P < 0.05, P < 0.01). i, NMDAR-related GSVA scores (Reactome and GO) in HPSNs across stages. j, GluN2B-endocytosis AUCell scores in HPSNs across stages. k,l, GluA2/NMDAR1 (k) and GAD1/NMDAR1 (l) immunofluorescence on sagittal brain sections, with mean-fluorescence quantification; merged images include DAPI and DBA (n = 3 mice per group). m, Neuroblast frequency among neuronal nuclei across stages. n,o, DCX and cleaved caspase-3 (CC3) immunofluorescence in dentate gyrus (n) and olfactory bulb (o) granule-cell layers, quantifying DCX+ cells in n and fluorescence intensity in o (3 mice per group). In panel n, *P < 0.05 for 0 versus 60 dpi. For c,d,g,j–o, one-way ANOVA with Tukey’s test. *P < 0.05, **P < 0.01, ***P < 0.001, ****P < 0.0001; ns, not significant.

To determine whether the neuropsychiatric abnormalities of cerebral toxoplasmosis are linked to excessive synaptic pruning, we performed Golgi staining on chronically infected brains and found significantly reduced dendritic spine density in cortical and hippocampal neurons (Fig. 4e). The complement genes *C1qa* and *C3* were both significantly upregulated after infection (Fig. 4f), and activated microglia colocalised with the excitatory postsynaptic marker PSD95 in the cortex (Fig. 4g). Quantified CD68–PSD95 colocalisation was essentially absent at 0 dpi and increased significantly at 12 and 60 dpi; at 60 dpi it displayed a marked spatial gradient, being minimal in cyst-free parenchyma, low around large cysts, and progressively higher around small cysts (< 40 μm diameter) and within microglial nodules (Fig. 4g), indicating that *T. gondii* infection drives excessive synaptic pruning through activated microglia, concentrated at sites of cyst-associated microglial aggregation.

To probe synaptic function, cell–cell communication analysis across all neurons showed that NMDAR signalling was broadly weakened after infection, whereas GABAergic inhibitory signalling was broadly enhanced (Fig. 4h); GSVA confirmed significant NMDAR-pathway downregulation in HPSNs (Fig. 4i). Consistent with inflammatory factors reducing synaptic activity via GluN2B endocytosis, the GluN2B-endocytosis gene set was significantly elevated at the transcriptional level (Fig. 4j), indicating reduced synaptic-membrane NMDAR availability. Immunofluorescence with quantitative analysis verified a significant, persistent decrease of NMDAR1 (GluN1) protein after infection (Fig. 4k,l), whereas the AMPA-receptor subunit GluA2 increased progressively from 0 to 60 dpi (Fig. 4k), and the GABAergic marker GAD1 decreased at 12 dpi before returning to baseline levels at 60 dpi (Fig. 4l). Thus, *T. gondii*-triggered neuroinflammation elicits a compensatory, protective remodeling: GABA-mediated inhibition is enhanced while glutamate-mediated excitation is reduced, limiting neuronal damage from hyperexcitability and calcium overload.

The adult mouse brain maintains a relatively stable neuronal population, yet *T. gondii* infection reduced neuronal numbers. Beyond structural injury and dysfunction of mature neurons, impaired adult neurogenesis may also contribute to these neuropsychiatric abnormalities. The number of neuroblasts, a neuronal subpopulation, declined significantly (Fig. 4m), and newborn neurons were markedly reduced in the dentate gyrus (Fig. 4n) and olfactory bulb (Fig. 4o) granule cell layers, demonstrating that *T. gondii* infection impairs adult neurogenesis.

### Spatial metabolomics reveals lipid reprogramming in the cyst microenvironment

Because the immune and atypical immune cells enriched in the CN are activated (Supplementary Fig. 3d), we asked whether the inflammatory microenvironment damages neurons by altering the local metabolite milieu. Immunofluorescence first revealed lipid peroxidation in the CN (Fig. 5a). Spatial metabolomics of chronically infected brain tissue, aligned with immunofluorescence on the same sections (Fig. 5b), compared the CN with nearby regions devoid of PV/cyst structures in the same brain area, identifying significantly altered metabolites (Supplementary Table 4). Parallel bulk metabolomics showed that the CN differential metabolites and enriched pathways were distinct from the bulk level (Fig. 5c–h): they were mainly enriched in glycerophosphoethanolamines, glycerophospholipids, lipids and lipid-like molecules, phosphatidylethanolamine (PE) and sphingolipids, with sphingolipid metabolism the most significantly downregulated integrated pathway (Fig. 5h).

**Fig. 5.**
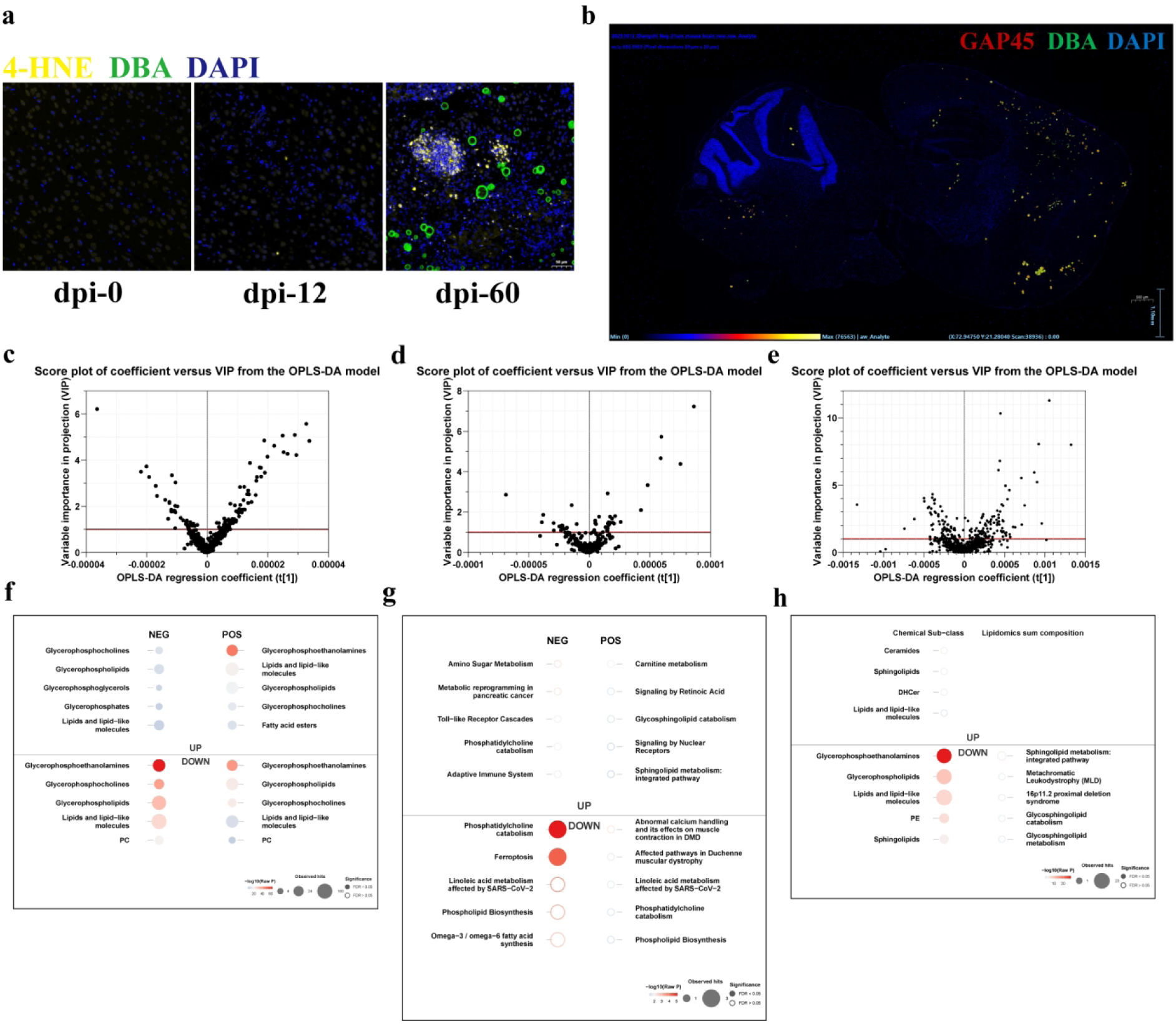
Spatial metabolomics reveals lipid reprogramming in the cyst microenvironment. a, Immunofluorescence of the lipid-peroxidation marker 4-HNE (yellow) with the cyst-wall marker DBA (green) and DAPI (blue) at 0, 12 and 60 dpi, showing 4-HNE accumulation around DBA^+^ cysts at 60 dpi, particularly in microglial nodule regions where cysts have ruptured. b, DESI-MSI of a chronically infected brain section (60 dpi) aligned with same-section immunofluorescence (GAP45, red; DBA, green; DAPI, blue); view in HDI software (ion image at m/z 830.5903, 20 μm × 20 μm pixels). Cyst neighbourhoods (GAP45^+^DBA^+^ foci) and matched cyst-free control regions in the same brain area were delineated as regions of interest for differential-metabolite analysis. c–e, OPLS-DA coefficient-versus-VIP (S-plot) score plots for the three metabolomics comparisons: bulk metabolomics, infected versus uninfected whole brain, negative ion mode (c); bulk metabolomics, positive ion mode (d); spatial metabolomics, cyst neighbourhood versus matched cyst-free regions, negative ion mode (e). Red lines mark the VIP = 1 threshold for differential metabolites (Supplementary Table 4; per-feature VIP scores and coefficients in Supplementary Table 7). f, Chemical sub-class enrichment of bulk differential metabolites, separated by ion mode (NEG, POS) and direction (up, down); dot colour, −log_10_(raw P); dot size, number of observed hits; filled dots, FDR < 0.05; open dots, FDR ≥ 0.05. g, Pathway enrichment of bulk differential metabolites (MetaboAnalyst enrichment library; one-sided hypergeometric test, Benjamini–Hochberg FDR < 0.05), by ion mode and direction as in f. h, Integrated chemical sub-class and lipidomics sum-composition enrichment of spatial cyst-neighbourhood differential metabolites, by direction (up, down); encodings as in f. Glycerophosphoethanolamines are the most significantly downregulated chemical sub-class, and sphingolipid metabolism ranks first among downregulated integrated pathways, in the cyst neighbourhood.

## Discussion

By integrating snRNA-seq with Stereo-seq, this study constructed a spatiotemporally resolved atlas of cerebral toxoplasmosis capturing both host cell populations and the parasite itself. The atlas portrays a dynamic equilibrium in the infected brain: cysts are continually disrupted, released bradyzoites reseed new cysts, and surrounding neural tissue accumulates damage. Three interconnected findings support this view. First, a cyst–neuron–microglia signalling axis recruits brain-resident immune cells to the cyst. Second, microglia disrupt cysts in a contact-dependent manner but paradoxically sustain persistent infection by releasing infectious bradyzoites. Third, synaptic, neurogenic and metabolic damage concentrates in the cyst microenvironment.

At the host level, infection remodeled a highly heterogeneous cellular landscape (18 major clusters, 80 subclusters). Among neurons, the persistent decline of HPSNs and the loss of multiple glutamatergic subtypes point to broadly compromised neuronal plasticity, providing a cellular basis for the cognitive and behavioural abnormalities of chronic infection. In contrast, highly active spiny neurons (HASNs) increased in the chronic phase; their high-metabolism, potentially high-transmitter-release signature may reflect host compensation or parasite manipulation (for example, interference with dopamine metabolism^38,39^) and awaits functional validation.

Astrocytes broadly transitioned from homeostatic states into diverse reactive states, including IEA and TRA (Supplementary Fig. 1c). IEA resembles neuroinflammatory reactive astrocytes and TRA resembles fibrotic reactive astrocytes, both of which have been reported in neurodegenerative disease and brain injury. IEA (A1-like astrocytes) lose synaptic-supporting and neurotrophic functions and secrete neurotoxic factors, consistent with a pro-neurotoxic microenvironment in the infected brain. Oligodendrocytes increased while OPCs declined in the chronic phase, suggesting that infection promotes OPC differentiation, possibly repairing demyelination but potentially depleting the OPC pool and compromising future remyelination. As first responders, microglia shifted from homeostatic states (HM_1/HM_2) towards IRM, PM and PIM states in the acute phase; in the chronic phase, homeostatic subsets partially recovered while activated states persisted, an equilibrium in which one subset limits tissue damage while another maintains surveillance of the cysts.

Peripheral immunity followed a clear recruitment sequence: neutrophils, mononuclear phagocytes and T cells infiltrated in the acute phase, whereas B cells and plasma cells appeared in large numbers only in the chronic phase. This sequence matches the time course of the antibody response and suggests that local humoral immunity plays an essential role in controlling chronic infection and preventing reactivation. Plasma cell colonisation and local antibody production in the brain may be a key mechanism limiting the escape of intracystic bradyzoites and neutralising extracellular parasites.

At the parasite level, six transcriptionally distinct subclusters coexisted in the brain. Systematic comparison with the bradyzoite subtypes of Ulu et al.^63^ revealed a broadly consistent architecture (Supplementary Fig. 4a–c): by Spearman correlation on 7,739 shared genes and diagnostic-marker analysis, subcluster 1 corresponded to Ulu Group B, the SRS22A^+^ subtype with reactivation potential; subcluster 2 corresponded to Group A, with a strong rhoptry program uncoupled from S/M-phase transcription; and subcluster 0 combined the cyst-wall maturation features of Groups C and D. Subcluster 5 recapitulated proliferative S/M features of Group E, although its highest global correlation was with Group D (ρ = 0.758). Subcluster 3 remained unclassified because its only candidate marker, GRA7, did not meet the FDR < 0.05 threshold (FDR = 0.05297). Subcluster 4 shared the SRS22A^+^/BAG1^lo^ state of Group B but additionally showed a microneme/AMA1 program and the highest AP2IX-9, making it a candidate reactivation-associated intermediate rather than a direct one-to-one match in the reference taxonomy. The two studies corresponded closely at the level of functional modules and gene families but overlapped weakly at individual marker genes (0–4 overlapping genes), consistent with robust gene programs whose individual composition varies between datasets. Our data support the SRS22A^+^/SRS22A^−^ framework and identify additional heterogeneity not resolved in the reference taxonomy. Subcluster 5 showed high ribosomal-protein expression and a distinct crowding-stress and antioxidant expression profile (Supplementary Fig. 4d), consistent with heterogeneous redox and metabolic adaptation across parasite states^64^.

Spatially, Stereo-seq linked cell identity to precise location, enabling dissection of the cyst neighbourhood. Stage classification of the structures themselves revealed how cyst composition evolves with growth: at 60 dpi, small structures were almost exclusively bradyzoite-dominant cysts, whereas the mixCyst fraction rose steeply with structure size, exceeding 90% of classifiable structures in the largest class (Supplementary Fig. 2f). Although replicating and tachyzoite-like parasites have previously been reported within tissue cysts^26^, our data provide, to our knowledge, the first spatially resolved evidence that newly formed chronic cysts are predominantly bradyzoite and that tachyzoite-or transitional-stage parasites emerge as cysts enlarge. Together with the SRS22A^+^ reactivation-potential subtype and the early reactivation intermediate resolved at single-cell level^63^, this size-graded mixing indicates that chronic cysts are not uniformly quiescent but progressively re-acquire tachyzoite-program activity as they grow, constituting a potential reservoir for reactivation and continual cyst turnover. Distance-based stratification (cCAC, cPAC, cDAC) revealed a cyst-centred immune microenvironment that changes in a gradient: microglia and mononuclear phagocytes increase toward the cyst, with the strongest immune activation around small cysts and stronger immune-evasion features in large cysts. Small PV/cyst structures are thus not immunologically silent: their neighbourhoods show the strongest immune activation, suggesting that young structures have an important role in sustaining chronic immune activation. Within this microenvironment, cyst-neighbourhood neurons strongly upregulate APP upon sensing the cyst or its secretions, and neuron-to-microglia communication is dominated by the App–Cd74 pair, strongest between HPSNs and IRMs at the cyst core. The APP–CD74 axis promotes pro-inflammatory^65^ and degenerative^66^ myeloid phenotypes in other disease contexts; we propose that here it constitutes a pathogen-driven neuroimmune injury axis: cyst-neighbourhood neurons recruit and activate microglia via APP–CD74, and the IRMs then recruit and activate cytotoxic T cells, which perforate the neuronal membrane to expose the intracellular cyst.

Microglia then act directly on the exposed cyst. N-SIM and TEM showed that microglia first adhere to the cyst surface with protrusions, then remodel their cytoskeleton into invasive protrusions that invade the cyst wall, disrupting its ultrastructure and releasing the bradyzoites within. On-chip single-cell analysis resolved this attack into three temporally ordered stages: environmental sensing and transcriptional activation, metabolic reprogramming, and effector execution combining protrusion extension, enzymatic digestion, oxidative killing, immune-cell recruitment and anti-apoptotic programs. This attack is thus a double-edged sword: intended to clear the pathogen, it instead releases bradyzoites and promotes secondary infection and de novo cyst formation; the reduced cyst numbers after microglial depletion corroborate this self-sustaining mechanism of chronic infection. Species differences are critical here: rodent cytotoxic lymphocytes lack granulysin, so after T cells disrupt the neuronal membrane, cyst destruction and killing of escaped parasites depend heavily on microglia and neighbouring phagocytes, which may partly explain the high susceptibility of mice to severe *T. gondii* infection. Human cytotoxic lymphocytes kill intracellular parasites directly via granulysin^37^, but whether they can penetrate the mature cyst wall is unknown. In immunosuppressed hosts such as AIDS patients, insufficient cytotoxic T cell recruitment allows escaped parasites to survive cyst disruption, reconvert to tachyzoites and proliferate unchecked, manifesting as severe toxoplasmic encephalitis; likewise, insufficient immune pressure at rupture sites in immunocompetent individuals would yield secondary cysts and maintain persistent latency. Microglial reactivity and nodule formation in the brains of people with HIV correlate with synaptic and dendritic damage, and nodules are more numerous in patients with neurological symptoms^67^, suggesting that cyst-disruption-associated microglial nodules may injure neurons; their association with *T. gondii* infection and cyst lysis in HIV-infected brains remains to be explored.

Neuronal injury in cerebral toxoplasmosis is therefore multifactorial: BBB damage manifests as progressive adhesion and residence of peripheral immune cells; activated astrocytes lose endfoot function, facilitating infiltration and amplifying parenchymal inflammation; and the cyst–neuron–microglia–T cell cascade amplifies local inflammation stepwise, with recurrent secondary cyst formation driving its chronicity. At the circuit level, infection weakens NMDAR signalling (with increased GluN2B endocytosis) while broadly enhancing GABAergic inhibition; upregulated complement (C1qa, C3) and the colocalisation of activated microglia with PSD95 point to excessive synaptic pruning that is strongest at microglial nodules and around small cysts and reduces dendritic spine density in cortex and hippocampus; and impaired adult neurogenesis in the dentate gyrus and reduced incorporation of newborn neurons into the olfactory bulb further compromise circuit repair^52,53,59^. Engulfment of postsynaptic material by microglia was significantly higher at microglial nodules and around small cysts than in cyst-free regions of the same brains (Fig. 4g); because nodules mark recent cyst-rupture sites and small cysts mark recent reseeding, this spatial coupling suggests that the cyst disruption– reseeding cycle is one driver of excessive synaptic pruning during chronic infection.

Spatial metabolomics revealed a metabolic dimension masked by bulk analysis. Four features stand out in the cyst neighbourhood: depletion of plasmalogens and polyunsaturated phosphatidylethanolamines (oxidative damage and membrane disintegration); reduction of glycosphingolipids and myelin-associated lipids (demyelination and white matter loss); accumulation of membrane-synthesis precursors alongside decreased mature membrane lipids (stalled synthesis with hyperactive breakdown); and elevation of pro-apoptotic sphingolipids such as ceramides (sphingolipid metabolic stress). We therefore propose a self-reinforcing vicious cycle in the cyst microenvironment: inflammation causes oxidative stress, oxidative stress drives lipid peroxidation, and lipid peroxidation in turn aggravates membrane damage and inflammation.

In summary, this study integrates spatiotemporal multi-omics with cellular and tissue-level analyses to examine interactions among cysts, neurons and microglia during *T. gondii* infection. The findings support a model in which microglia participate in cyst disruption and secondary cyst formation, while changes in neuronal signalling, neurogenesis and lipid metabolism are associated with brain injury. The resulting atlas provides a resource for investigating host–parasite interactions in the infected brain. Further work is needed to define the causal relationships between cyst turnover and specific neuronal injuries and to determine whether modulating microglial activity can limit pathology without compromising parasite control.

## Methods

### Mice and parasite strains

All animal experiments were conducted in accordance with the Regulations for the Administration of Affairs Concerning Experimental Animals (China) and were approved by the Southern Medical University Animal Ethics Committee (permit no. SMUL202308003). Wild-type 129S2/SvPasCrl mice (6-week-old males, 18–22 g) were bred and housed in the institutional Laboratory Animal Center under specific-pathogen-free conditions (20–26 °C, 40–70% relative humidity, 12-h light/12-h dark cycle), with food and water ad libitum. *Toxoplasma gondii* ME49 strain (type II) was maintained by serial oral passage of tissue cysts in 129S2/SvPasCrl mice. Tissue cysts were purified from the brains of mice chronically infected for 2 months by density-gradient centrifugation over lymphocyte separation medium (Histopaque-1077, Sigma-Aldrich; density 1.077 g/mL)^68^, adapting the method of Cornelissen *et al.*^69^. Briefly, four brains were pooled, homogenised in a glass tissue homogeniser, and brought to 30 mL with PBS. Each 15 mL of homogenate was layered over 20 mL of separation medium and centrifuged at ≈250 × g for 30 min at 25 °C in a swinging-bucket rotor. A low g force was chosen because young cysts are vulnerable to rupture at high g forces, with consequent release of bradyzoites, and intact cysts were required for downstream experiments. The bottom ≈5 mL of each tube, containing the cysts, was collected, filtered through sterile gauze, and pelleted at ≈500 × g for 10 min. Cysts were washed once in PBS and counted under a light microscope before use.

### Primary cell culture

All primary cells were obtained from 129S2/SvPasCrl mice housed under specific pathogen-free (SPF) conditions. Complete medium consisted of DMEM (Gibco, C11995500BT) supplemented with 10% FBS (Gibco, 10099141c). Brains from 3-day-old (P3) mice were isolated, freed of blood vessels and meninges, minced, digested and mechanically dissociated into single-cell suspensions, which were seeded in culture plates and maintained at 37 °C in a 5% CO_2_ incubator.

For microglia, after overnight culture the medium was replaced with complete medium and fully changed every 48–72 h. After approximately one week, microglia were detached from the underlying astrocyte layer by gentle pipetting, collected by centrifugation (300 × g, 5 min) and resuspended for use.

For astrocytes, the medium was replaced with complete medium after overnight culture, switched 48–72 h later to complete medium containing 1 μM PLX5622 (to deplete contaminating microglia) for 48–72 h, and then returned to PLX5622-free complete medium for a further 48–72 h before experiments.

### Infection model and omics sample preparation

Mice were infected orally with five ME49 cysts each. Brains were collected at 0 (uninfected), 12 (acute) and 60 (chronic) days post-infection, with three mice per time point as biological replicates. For snRNA-seq, whole brains were rinsed twice in PBS, one hemisphere was rapidly dissected along the sagittal midline and snap-frozen in liquid nitrogen, and nuclei were isolated by Dounce homogenisation followed by iodixanol density-gradient centrifugation. For Stereo-seq, brains were embedded in OCT, snap-frozen, sectioned at 10 μm and mounted onto 1 × 2 cm Stereo-seq chips (BGI Research, Shenzhen). Adjacent sections were processed for histology (H&E) to confirm parasite invasion and inflammation.

### snRNA-seq library preparation and sequencing

Nuclear suspensions (20,000 nuclei per sample) were loaded onto a 10x Genomics Chromium instrument, and libraries were constructed using the Chromium Next GEM Single Cell 3’ Reagent Kit v3.1 according to the manufacturer’s instructions. Briefly, gel bead-in-emulsions (GEMs) were generated; after reverse transcription, barcoded full-length cDNA was amplified by PCR (11 cycles), and libraries were prepared by end repair, A-tailing, adaptor ligation and sample-index PCR. Libraries were quantified by Qubit and sequenced on an Illumina NovaSeq X Plus platform (paired-end 150 bp) to a depth of 21,000–52,000 reads per nucleus (mean, 33,000 reads per nucleus).

### Stereo-seq library preparation and sequencing

Stereo-seq was performed following the manufacturer’s protocol (BGI). Frozen sections (10 μm) were adhered to tissue-optimisation and DNB chips, fixed with pre-cooled methanol, and stained with Qubit ssDNA dye to record nuclei positions by fluorescence microscopy. Tissue permeabilisation time was optimised on the optimisation chip (optimal time, 12 min). mRNA captured by poly-T probes on the chip was reverse-transcribed in situ, and cDNA was amplified, fragmented to 200– 300 bp and used to construct libraries with P5/P7 adaptors and dual indices. Libraries were sequenced on an MGI DNBSEQ-Tx platform in PE50+100 mode: read 1 contains 25-bp coordinate identity (CID) and 10-bp molecular identifier (MID) information, and read 2 contains the cDNA sequence.

### Stereo-cell chip co-culture and sequencing

Primary microglia were co-cultured with purified ME49 cysts on Stereo-cell chips (BGI) for 6 h at 37 °C, 5% CO_2_, following the established Stereo-cell workflow^70^. After sequencing, the raw reads were processed using the STOmics Analysis Workflow (SAW, BGI) for quality control, alignment to the reference genome and gene expression quantification. Cells adjacent to cysts were manually gated on chip images to define cyst-neighbourhood microglia (CNM) and non-neighbourhood microglia (NNM). Differentially expressed genes between CNM and NNM were identified with the Wilcoxon rank-sum test in Scanpy (rank_genes_groups), with thresholds of |log_2_FC| > 0.5 and *P* < 0.05 (Supplementary Table 3).

### snRNA-seq data processing, clustering and sub-clustering

Raw reads were aligned to a combined mouse (mm_release112) and *T. gondii* ME49 (ToxoDB-68_TgondiiME49) reference using Cell Ranger with default parameters. Downstream analysis used Seurat (v5.1.0) in R (v4.3.2). Cells/nuclei with < 400 genes, > 6,500 genes, UMI > 27,000, or > 5% mitochondrial reads were excluded; doublets were removed with DoubletFinder. Data were normalised (LogNormalize), integrated across samples with Harmony, and clustered using Louvain at resolution 0.5. Major cell populations were annotated by canonical markers shown in Fig. 1d, and each major population was sub-clustered independently to yield 80 subpopulations. Parasite cells were analysed analogously; pseudotime trajectories were inferred with Monocle2. Differential expression analysis used MAST (Model-based Analysis of Single-cell Transcriptomics) with thresholds of |log_2_FC| > 1 and *P* ≤ 0.05.

### Parasite life-stage classification in single-cell data

Parasite cells were scored for two stage-specific gene modules using AUCell: a tachyzoite module (SAG1, SAG2A, SRS1, SRS2, LDH1, ENO2, AMA1) and a bradyzoite module (BAG1, LDH2, SRS35A, MAG1, ENO1, CST1, SRS13; gene identifiers in Supplementary Table 5). Of 2,188 parasites (28 at 12 dpi; 2,160 at 60 dpi), 42 cells with zero counts for all 14 module genes were excluded from threshold estimation and downstream classification, leaving 2,146 cells. Classification thresholds were defined from the empirical score distributions: θ_B_ = 0.18 corresponded to the local minimum of the bradyzoite-score distribution separating the basal low-signal shoulder from the main bradyzoite-active population, and θ_T_ = 0.15 was set at the inflection of the tachyzoite-score distribution discriminating the low-level basal (ambient-RNA-consistent) shoulder from the sparse tachyzoite-active tail (Supplementary Fig. 1o,p). Cells with T ≥ θ_T_ and B < θ_B_ were classified as Tachyzoite; T < θ_T_ and B ≥ θ_B_ as Bradyzoite; T ≥ θ_T_ and B ≥ θ_B_ as Transitional (dual-positive); and T < θ_T_ and B < θ_B_ as Undetermined. An alternative rule additionally requiring ≥ 2 of 7 module genes with non-zero counts for a module to be considered active yielded concordant assignments (Supplementary Fig. 1q); per-cell module scores and both classification calls are provided in Supplementary Table 1. Robustness was assessed by ±10% and ±20% perturbation of both thresholds (16 combinations; agreement 88.8–98.8%; Supplementary Fig. 1r and Supplementary Table 6). Parasite recovery at 12 dpi was limited (n = 28) for technical reasons; 12-dpi stage inference therefore relies primarily on spatial data, with single-cell results presented as concordant evidence.

### Spatial deconvolution, cyst identification and cyst-neighbourhood analysis

Stereo-seq bins (cell bin) were deconvoluted against the snRNA-seq reference to map the 18 host populations in space, using Tangram for major populations and non-neuronal subclusters and Stereoscope for neuronal subclusters. PV/cyst structures were identified from the on-chip ssDNA signal and parasite transcript signal. Stage classification of parasite foci in the spatial data was performed as follows. Parasite foci (candidate parasitophorous vacuoles or tissue cysts) were manually delineated on Stereo-seq sections from infected mouse brains (140 foci at 12 dpi; 300 at 60 dpi). Nineteen foci lacking detectable parasite UMIs were excluded, leaving 421 foci. For each focus i and parasite gene g, expression was normalised within the parasite compartment as logCPM(g, i) = log_2_(1 + 10⁶ × X(g,i) / Σg X(g,i)), where the denominator sums over parasite genes only; host transcripts, focus area and housekeeping genes were not used as denominators. Module scores T and B were defined as the mean logCPM of the tachyzoite and bradyzoite panel genes, respectively. Thresholds θ_T_ = θ_B_ = 0.30 were set at the local minimum following the zero-inflated spike in each score distribution (Supplementary Fig. 2b). Foci were classified as parasitophorous vacuoles (PV; T ≥ θ_T_, B < θ_B_), tissue cysts (Cyst; T < θ_T_, B ≥ θ_B_) or mixed-stage cysts (mixCyst; T ≥ θ_T_, B ≥ θ_B_). Foci with both scores below threshold, or in which a supra-threshold module score was supported by fewer than two detected panel genes, were conservatively designated Unresolved. Class composition was summarised per time point with individual mice as biological replicates; per-focus scores and classifications are provided in Supplementary Table 2. All PV/cyst structures were included in size-class and neighbourhood analyses. Cross-sectional areas were computed and classified by K-means (k = 4) into four size classes (sCs, 0–1,100 μm²; mCs, 1,100–2,550 μm²; lCs, 2,550–5,000 μm²; xCs, 5,000–14,000 μm²). Neighbourhoods were defined by distance to each structure: cCAC (inside the structure to 15 μm beyond its border), cPAC (15–45 μm), cDAC (45–75 μm); remaining 60-dpi cells served as structure-non-associated background (cBCs), and 0-dpi cells as uninfected controls (UCC). Cell-type enrichment per layer was tested against cBC by Wilcoxon test. BBB adhesion of peripheral immune cells was quantified by a proximity score to endothelial cells (threshold > 0.414; Supplementary Fig. 3c). Ligand–receptor interactions between cell populations were inferred with CellChat 1.6.1, run separately per infection stage and per neighbourhood layer; communication probabilities and pathway-level signalling were computed with default parameters.

### AUCell, GSVA, GO and KEGG analyses

Gene-set scoring used AUCell (v1.28.0) and GSVA (v1.30.0) with gene sets from MSigDB (v7.4), Gene Ontology (GO) and the Kyoto Encyclopedia of Genes and Genomes (KEGG). GO and KEGG enrichment of DEGs was performed with clusterProfiler 4.14.0 using a hypergeometric test; terms with adjusted *P* < 0.05 (Benjamini–Hochberg) were considered significant. The TOP23/TOP56/TOP72 gene sets were derived by ranking CNM-versus-NNM DEGs by significance (Supplementary Table 3).

### Spatial and bulk metabolomics

For spatial metabolomics, frozen brain sections (10 μm) from mice chronically infected for 60 days were analysed by desorption electrospray ionisation mass spectrometry imaging (DESI-MSI) on a Waters SELECT SERIES™ MRT (multi-reflecting time-of-flight) system in negative ion mode at a spatial resolution of 20 μm; immunofluorescence was performed on the same sections for alignment. Cyst neighbourhoods and matched cyst-free regions of interest were delineated using HDI v1.8. For bulk metabolomics, whole-brain homogenates from infected and uninfected mice (n = 6 per group) were extracted with isopropanol (IPA) and analysed by ultra-high-performance liquid chromatography–quadrupole time-of-flight mass spectrometry (Waters ACQUITY Premier System) equipped with an ACQUITY Premier BEH C18 1.7 μm VanGuard FIT column (2.1 × 100 mm). Electrospray ionisation (ESI) was performed in both positive and negative ion modes. The mobile phase consisted of phase A (60% acetonitrile / 40% water) and phase B (90% isopropanol / 10% acetonitrile), both containing 10 mM ammonium formate and 0.1% formic acid. The injection volume was 5 μL, the column temperature was 60 °C, and the autosampler was maintained at 10 °C. The mass axis was calibrated using a sodium formate solution (0.5 mmol/L). Leucine enkephalin (0.2 μg/mL) was used as the lock mass for detector calibration and as an internal standard. The mobile-phase gradient and mass spectrometry parameters are described in Supplementary Methods 1. Metabolites were identified against the LIPID MAPS v2013.11.21.01, HMDB 2017v and METLIN MS/MS Library 2019 databases. Differential metabolites were defined by variable importance in projection (VIP) > 1 (per-feature VIP scores and OPLS-DA coefficients in Supplementary Table 7). Pathway enrichment was performed using the MetaboAnalyst enrichment library via MetaboAnalystR 4.3.0. Enrichment *P* values were calculated using a one-sided hypergeometric test. Multiple-testing correction was performed using the Benjamini–Hochberg procedure, and an FDR below 0.05 was considered statistically significant. Holm-adjusted *P* values were also retained in the source result tables but were not used as the primary significance criterion.

### Immunofluorescence

Brains were fixed in 4% PFA and sectioned at 5 μm. Sections were permeabilised/blocked in Triton X-100/5% BSA and incubated overnight at 4 °C with primary antibodies or fluorophore-conjugated reagents (anti-APP, Abcam, ab32136, 1:200; anti-CD74, Santa Cruz, sc-20062, 1:200; DBA-FITC, Invitrogen™, L32474, 1:400; anti-IBA1, Abcam, ab283319, 1:100; anti-IBA1, Abcam, ab178847, 1:200; iFluor™ 594 phalloidin, YEASEN, 40786ES75, 1 µg/mL; anti-GFAP, Abcam, ab279290, 1:100; anti-TgGAP45, generated in-house, 1:100; anti-CD68, Servicebio, GB153109, 1:300; anti-PSD95, Servicebio, GB150080, 1:50; anti-NMDAR1, Servicebio, GB11144, 1:100; anti-GluA2, Servicebio, GB111578, 1:400; anti-GAD1, Servicebio, GB150717, 1:1000; anti-Doublecortin, Abcam, ab18723, 1:200; anti-Cleaved Caspase-3, Affinity, BF0711, 1:200; anti-4-HNE, Servicebio, GB150073, 1:500; anti-MAP2, Servicebio, GB155559, 1:300), then with fluorophore-conjugated secondary antibodies (647-conjugated goat anti-rabbit IgG, Baiqiandu, C4121, 1:200; CY3-conjugated goat anti-mouse IgG, Baiqiandu, C4112, 1:200; Alexa Fluor 594-conjugated goat anti-mouse IgG (H+L), Invitrogen™, A35510, 1:200; Alexa Fluor 594-conjugated goat anti-rabbit IgG (H+L), Invitrogen™, A32740, 1:200; Alexa Fluor 488-conjugated goat anti-mouse IgG (H+L), Invitrogen™, A11001, 1:200) for 1 hour. Nuclei were counterstained with DAPI. Fluorescence images were acquired and analysed with a microscope (Nikon) and slide scanners (Olympus VS200; 3DHISTECH CaseViewer, Pannoramic SCAN).

### Golgi staining

Golgi staining was performed with Golgi staining solution (Servicebio, G1069-1) and Golgi fixative solution (Servicebio, G1069) on 4 mice per group according to the manufacturer’s instructions. 60 μm sections were cut; well-impregnated neurons per region (cortex, hippocampus) per mouse were imaged with a digital pathology slide scanner (Servicebio, LG-S80), and dendritic spines were counted on complete neurons along dendritic branches over a 30–90 μm length range. The branch length was measured and the number of spines within that length was counted; spine density was expressed as the number of spines per 10 μm, calculated as (number of spines / dendritic length) × 10, using Image-Pro Plus 6.0. Analysis was performed blinded to group.

### N-SIM super-resolution microscopy

Co-cultures of primary microglia and purified cysts were fixed at 24 hours with 4% PFA and imaged on a Nikon N-SIM super-resolution microscope system. 3D images were processed with Imaris 8.1 software (Nikon).

### Transmission electron microscopy

Co-culture samples were fixed in 2.5% glutaraldehyde in buffer, post-fixed in 1% osmium tetroxide, dehydrated through graded ethanol, embedded in resin and sectioned at 70 nm (Leica UC7). Sections were stained with uranyl acetate and lead citrate and imaged on a Hitachi H-7500 transmission electron microscope at 60 kV.

### PLX5622 microglia depletion

From 1 month post-infection, mice received PLX5622 (1200 ppm in AIN-76A diet) or PLX5622-free AIN-76A diet as control for 1 month (n = 5 treated; n = 3 untreated). Because cyst diameter spans multiple adjacent sections, a single complete sagittal section per mouse (olfactory bulb through cerebellum and medulla) was used for whole-section cyst counting. Cysts were enumerated and diameters measured on H&E-stained sections by an investigator blinded to treatment.

### Statistics and reproducibility

The number (n) of replicates/animals used per group is described in each figure legend and represents biological replicates. Statistical analyses were performed in R (v4.3.2) and GraphPad Prism 10. Ordinary one-way ANOVA with Tukey’s multiple comparisons test was used to evaluate experiments containing more than two groups; comparisons between two groups used the unpaired t-test with Welch’s correction or the Mann–Whitney test, as indicated in the figure legends, and data are presented as mean ± s.e.m. unless otherwise stated. Raw *P* values were used for the Stereo-cell CNM-versus-NNM DEG screen without Benjamini–Hochberg correction; adjusted *P* values from Benjamini–Hochberg are identified explicitly where used. No statistical methods were used to predetermine sample sizes; sample sizes were chosen on the basis of prior experience with this infection model and are similar to those reported in previous publications. Animals were randomly allocated to experimental groups where applicable. *P* < 0.05 was considered significant; exact *P* values are reported in the figure legends. Investigators were blinded to group allocation during quantification steps.

## Data availability

The snRNA-seq and Stereo-seq data generated in this study have been deposited in the Genome Sequence Archive (GSA) under accession numbers CRA048999 (https://ngdc.cncb.ac.cn/gsa/browse/CRA048999) and CRA049065 (https://ngdc.cncb.ac.cn/gsa/browse/CRA049065), respectively.

## Code availability

Custom code used for the analyses in this study is available at https://github.com/zhangchismu/SC-ST-code.

## Acknowledgements

This research was supported by Key project of National Natural Science Foundation of China (82330072), National Natural Science Foundation of China (82272364), and Guangdong Provincial Natural Science Foundation (2023A1515011733, 2024A1515011327) to HJP.

## Author contributions

Conceptualization: Chi Zhang. Investigation: Chi Zhang, Yingying Lu, Weiling Wu, Wenjun Huang and Xinyue Liao. Data analysis: Chi Zhang, Yingying Lu, Zhihao Zhou, Shuyu Zheng and Xiangyi He. Writing – original draft: Chi Zhang. Writing – review & editing: Hongjuan Peng. Supervision: Hongjuan Peng, Jiating Chen, Min Chen, Weihao Zou and Lijuan Zhou. Funding acquisition: Hongjuan Peng, Chi Zhang and Jiating Chen.

## Additional information

Supplementary information: Supplementary Figs. 1–4, Supplementary Tables 1–7, Supplementary Note 1 (cell-subcluster annotation) and Supplementary Methods 1 (LC–MS mobile-phase gradient and mass-spectrometry parameters) accompany this article.

---

**Supplementary Fig. 1.**
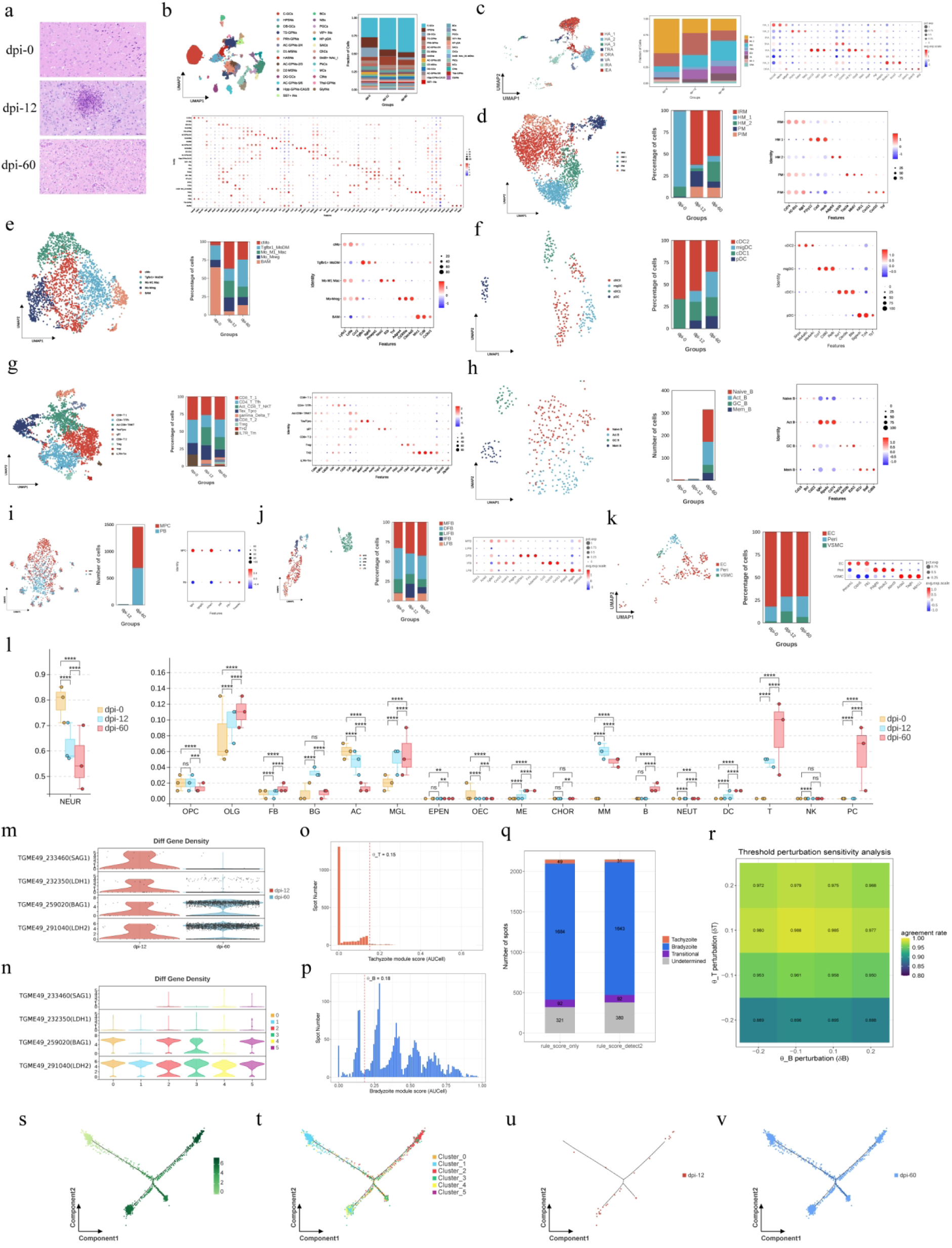
Host cell subclustering, parasite marker-gene expression and stage-classification quality controls. a, H&E staining of brain sections at 0, 12 and 60 dpi. b–k, UMAP, stage composition and marker-gene dot plots for major host populations: neurons (b), astrocytes (c), microglia (d), monocytes/macrophages (e), dendritic cells (f), T cells (g), B cells (h), plasma cells (i), fibroblasts (j), and mural/endothelial cells (k). Annotations are in Supplementary Note 1. l, Major cell-population frequencies across stages (n = 3 mice per time point; one-way ANOVA with Tukey’s test; ns, not significant; *P < 0.05, **P < 0.01, ***P < 0.001, ****P < 0.0001). m,n, SAG1, LDH1, BAG1 and LDH2 expression by infection stage (m) and parasite subcluster (n). o,p, Tachyzoite (o) and bradyzoite (p) AUCell-score distributions for 2,146 parasites. Dashed lines mark θT = 0.15 and θB = 0.18; threshold determination is described in Methods. q, Stage-class counts using score thresholds alone or with an additional requirement of ≥ 2 detected genes per active module (score-only versus detection rule: Bradyzoite, 1,684 versus 1,643; Tachyzoite, 49 versus 31; Transitional, 92 versus 92; Undetermined, 321 versus 380; Supplementary Table 1). r, Agreement of stage assignments across 16 combinations of ±10% and ±20% threshold perturbations (agreement, 88.8–98.8%; Supplementary Table 6). s–v, Monocle 2 pseudotime trajectories of six parasite subclusters, coloured by pseudotime (s), subcluster (t), and cells from 12 dpi (u) and 60 dpi (v).

**Supplementary Fig. 2.**
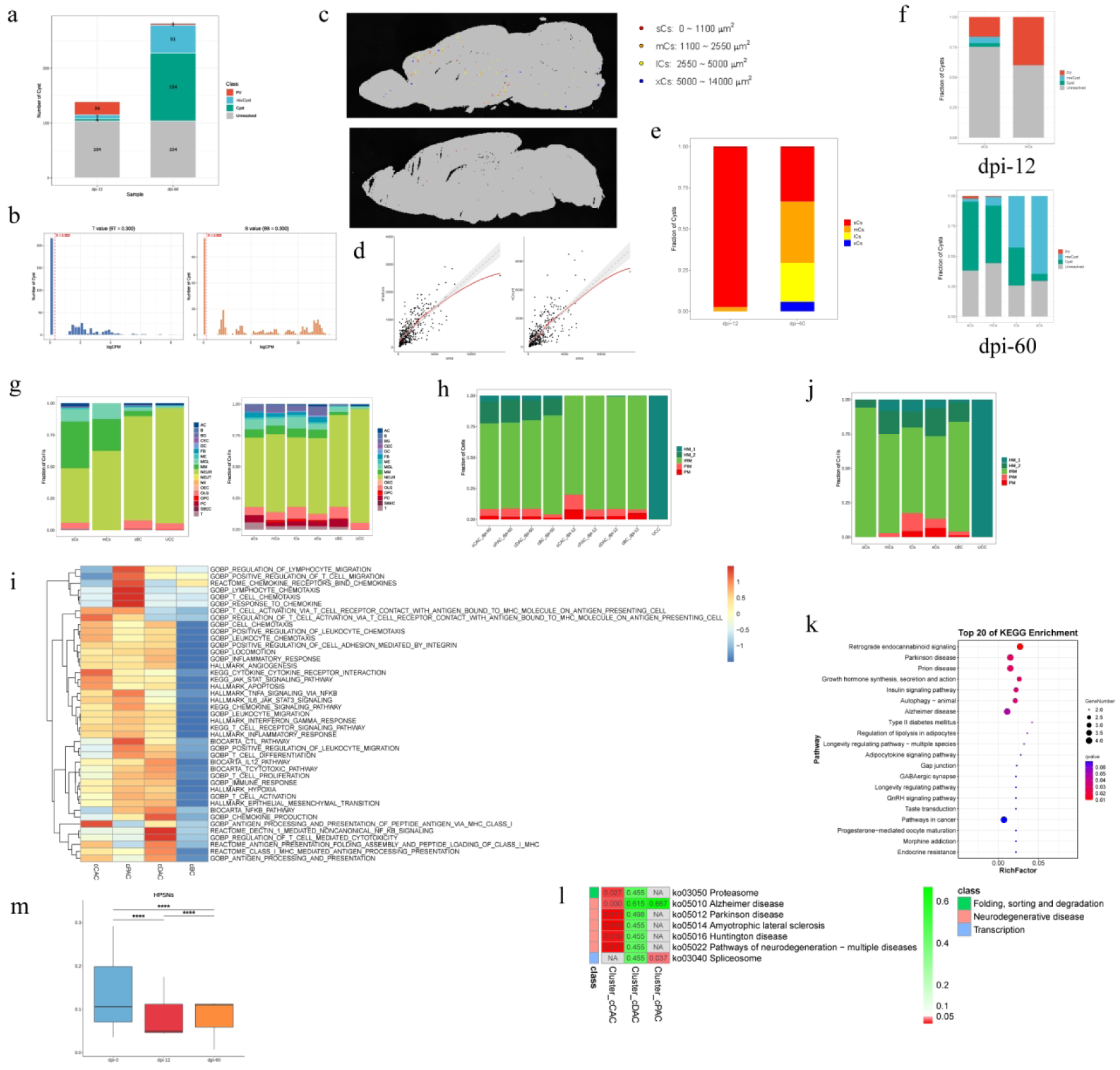
Stage and size classification of PV/cyst structures and cyst-neighbourhood composition. a, Stage classification of parasite foci. Of 440 delineated foci, 19 without parasite UMIs were excluded, leaving 421 (139 at 12 dpi; 282 at 60 dpi). Tachyzoite (T) and bradyzoite (B) logCPM module scores (θ_T_ = θ_B_ = 0.30) defined PV (T^+^B^−^), Cyst (T^−^ B^+^), mixCyst (T^+^B^+^) and Unresolved. At 12 dpi, counts were PV 24, mixCyst 7, Cyst 4 and Unresolved 104; at 60 dpi, PV 3, mixCyst 51, Cyst 124 and Unresolved 104. PVs comprised 24 of 35 classifiable 12-dpi foci, and cysts (Cyst plus mixCyst) 175 of 178 classifiable 60-dpi foci (Supplementary Table 2). b, T and B module-score distributions across 421 foci; dashed lines mark θ_T_ = θ_B_ = 0.30. c, Spatial distributions of size classes (sCs, mCs, lCs, xCs) at 12 and 60 dpi. d, Detected transcripts (nCount) and genes (nFeature) per structure versus cross-sectional area, with linear regression fits (nCount–area: R = 0.722, P = 5.84 × 10^-^⁶²; nFeature–area: R = 0.745, P = 6.47 ×10^-^⁶⁸). e,f, Size-class composition by infection stage (e) and stage composition within each size class (f). At 60 dpi, mixCyst accounted for 4%, 13%, 58% and 92% of classifiable sCs, mCs, lCs and xCs, respectively. g, Host cell-type composition of CN by size class, with cBC and UCC as references. h, Microglial subcluster composition across CN layers at 12 and 60 dpi. i, GSVA of cytotoxic CD8^+^ T-cell chemotaxis, activation and differentiation pathways in CN-associated IRMs across layers and cBC. j, Microglial subcluster composition by structure size class. k,l, KEGG enrichment for DEGs from microglia associated with large structures (k, top 20 terms) and across CN layers (l); dot size, gene number; colour, q value. m, HPSN frequency across infection stages (n = 3 mice per time point; one-way ANOVA with Tukey’s test; P < 0.001, P < 0.0001).

**Supplementary Fig. 3.**
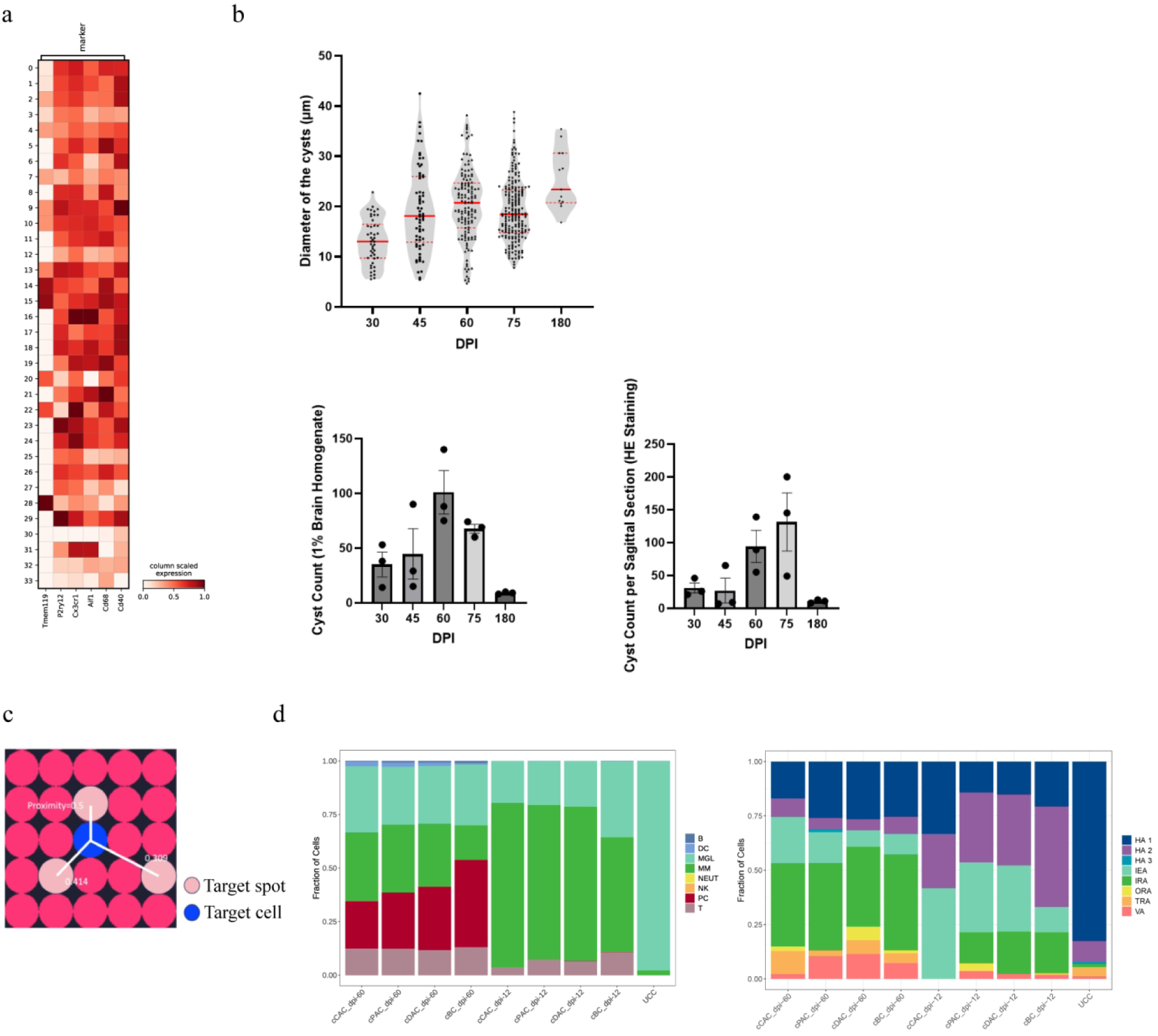
Stereo-cell recovery controls, cyst-burden kinetics, proximity-score schematic and CN composition. a, Marker-expression heatmap (column-scaled) of cells recovered from Stereo-cell chips after the 6-h co-culture (microglial markers Tmem119, P2ry12, Cx3cr1, Aif1, Cd68, Cd40); rows are the recovered on-chip cell clusters (0–33), confirming that nearly all recovered cells are primary microglia. b, Cyst-burden kinetics in the 129S2/ME49 oral-infection model: cyst diameter (top; violins with individual cysts as dots; red solid lines, medians; red dashed lines, quartiles) and cyst counts (bottom left, per 1% brain homogenate; bottom right, per sagittal H&E section; bars show mean ± s.e.m., individual mice shown as dots) at 30, 45, 60, 75 and 180 dpi (n = 3 mice per time point). c, Schematic of the proximity score used to define BBB adhesion: the score between a query spot and target spots is computed from their spatial relationship (illustrated values 0.5, 0.414 and 0.309); peripheral immune cells with proximity to endothelial cells > 0.414 were defined as BBB-adherent. d, Composition of immune cell populations (left) and astrocyte activation states (HA1, HA2, HA3, IRA, TRA, IEA, VA, ORA; right) across CN layers at 12 and 60 dpi, with cBC and UCC as references, showing activation of CN-enriched immune and atypical immune cells.

**Supplementary Fig. 4.**
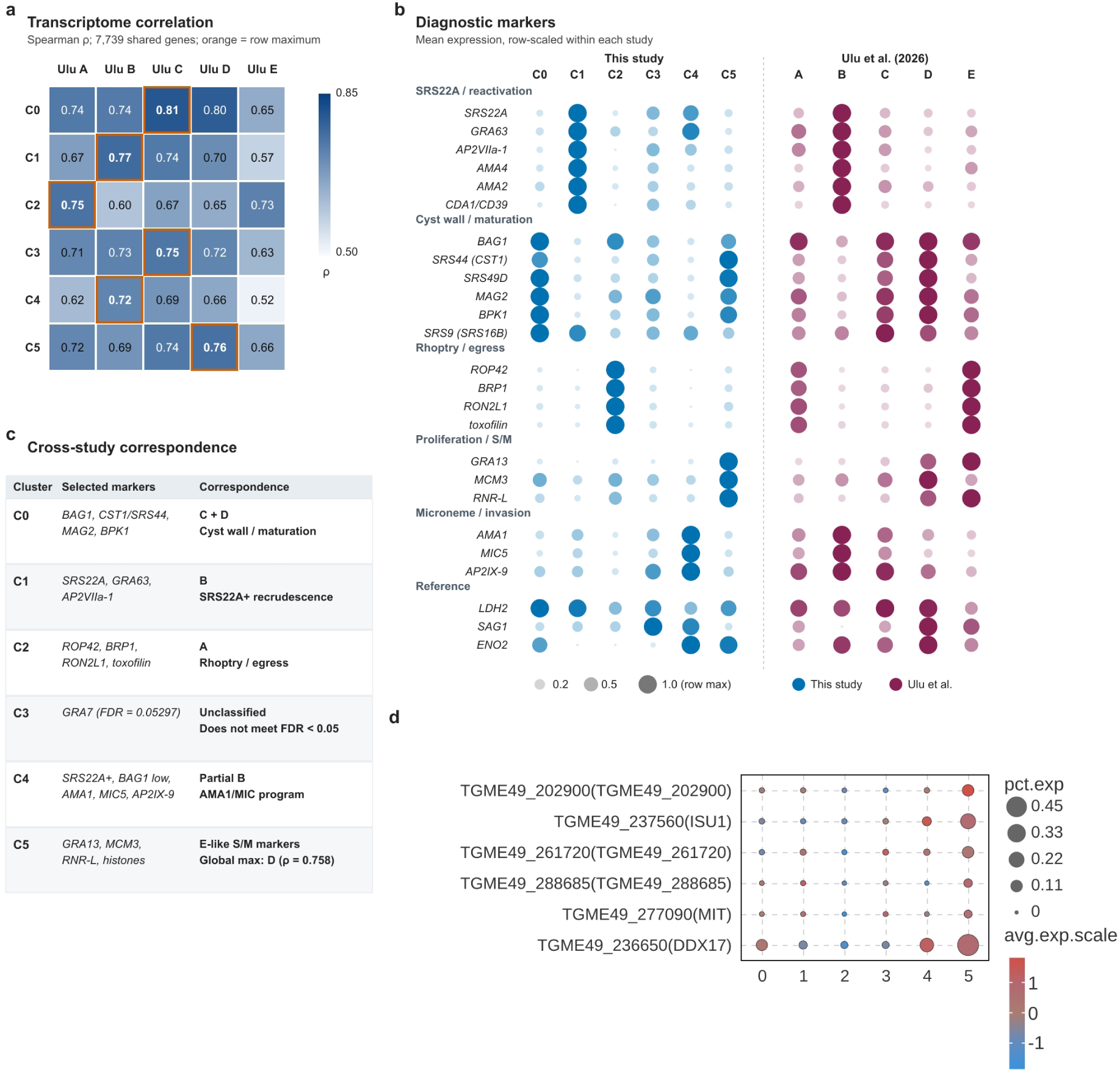
Cross-study comparison of bradyzoite subtypes and stress/adaptation marker expression. a, Spearman correlation of mean expression over 7,739 shared genes between six parasite subclusters (C0–C5) and five in vivo bradyzoite groups (A–E) from Ulu et al. Orange frames denote the highest correlation in each row. b, Mean expression of 25 diagnostic markers, scaled to the maximum for each gene within each study. Circle size and opacity denote the fraction of the row maximum. Modules group markers by biological program. c, Cross-study correspondence based on global correlations and diagnostic markers. C5 shows E-like S/M-marker expression despite its highest global correlation with Ulu D (ρ = 0.758); C4 partially overlaps with Ulu B but adds a microneme/AMA1 program; C3 is unclassified because its only candidate marker, GRA7, does not meet FDR < 0.05 (FDR = 0.05297). d, Dot plot of crowding-stress- and antioxidant-related transcripts (TgPRO (TGME49_202900), TgISU1 (TGME49_237560), TGME49_261720 (ZIP-family metal cation transporter), HscB (TGME49_288685; Fe–S protein assembly co-chaperone), TgMIT (TGME49_277090) and TgDDX17 (TGME49_236650)) across parasite subclusters 0–5. Dot size indicates the percentage of expressing cells (pct.exp); colour indicates scaled mean expression (avg.exp.scale).

## Supplementary Tables

Full data are provided in the accompanying Excel files. Each workbook includes a Table legend worksheet containing its title and description.

**Supplementary Table 1 | Per-cell tachyzoite/bradyzoite module scores and stage classification of intracerebral parasites.** Sheet 1 (Per-cell module scores): for each of the 2,146 analysed parasite cells: cell ID, tachyzoite- and bradyzoite-module AUCell scores, sample, subcluster (0–5), time point (12 or 60 dpi), and the stage calls under the score-only rule (rule_score_only) and the rule additionally requiring ≥ 2 detected module genes (rule_score_detect2). Sheet 2 (Classification rule counts): class counts under each rule. Data correspond to Fig. 1g, h and Supplementary Fig. 1m–r.

*Excel file: Suppl_Table1_per_cell_classification.xlsx*

**Supplementary Table 2 | Per-focus module scores and stage classification of spatial PV/cyst structures.** Sheet 1 (Per-focus classification): for each of the 421 analysed parasite foci: focus ID, tachyzoite (T) and bradyzoite (B) module scores (mean within-parasite logCPM), time point, cross-sectional area, numbers of detected tachyzoite (nzT) and bradyzoite (nzB) panel genes, and the assigned class (PV, mixCyst, Cyst, Unresolved). Sheet 2 (Focus class counts): class counts per time point. Data correspond to Supplementary Fig. 2a.

*Excel file: Suppl_Table2_per_focus_classification.xlsx*

**Supplementary Table 3 | expressedDifferentially expressed genes between cyst-neighbourhood microglia (CNM) and non-neighbourhood microglia (NNM).** The worksheet (CNM vs NNM DEGs) lists genes detected as differentially expressed between CNM and NNM recovered from Stereo-cell chips (Wilcoxon rank-sum test; |log2 fold change| > 0.5 and P < 0.05). Columns: gene symbol, Wilcoxon score, log fold change, raw P value, Benjamini–Hochberg-adjusted P value, absolute log2 fold change and Ensembl gene ID. Data correspond to Fig. 3h, i.

*Excel file: Suppl_Table3_CNM_vs_NNM_DEG.xlsx*

**Supplementary Table 4 | Differential metabolites from bulk and spatial metabolomics.**Metabolites with VIP > 1. Worksheets: NEG UP, bulk negative-ion upregulated; NEG DOWN, bulk negative-ion downregulated; POS UP, bulk positive-ion upregulated; POS DOWN, bulk positive-ion downregulated; SM UP, spatial cyst-neighbourhood upregulated; SM DOWN, spatial cyst-neighbourhood downregulated. Columns are Compound ID (HMDB/LIPID MAPS where available) and Description. Data correspond to Fig. 5c–h; per-feature VIP scores and coefficients are provided in Supplementary Table 7.

*Excel file: Suppl_Table4_differential_metabolites.xlsx*

**Supplementary Table 5 | Tachyzoite and bradyzoite stage-module genes.** The single worksheet (Stage module genes) lists the 14 genes used for tachyzoite/bradyzoite stage classification (7 genes per module). Columns: module assignment, gene symbol, synonyms, ToxoDB gene ID (ME49) and protein description. These gene panels were used for AUCell-based classification of parasite single cells and for spatial PV/cyst focus classification as described in Methods.

*Excel file: Suppl_Table5_stage_module_genes.xlsx*

**Supplementary Table 6 | Sensitivity of parasite stage classification to threshold perturbations.** The worksheet (Threshold sensitivity) reports 16 combinations of relative perturbations (dT and dB: −0.2, −0.1, +0.1 and +0.2) applied to the adopted tachyzoite and bradyzoite thresholds (thetaT and thetaB), and the resulting agreement rate (agree_rate) with the reference classification. The adopted thresholds were thetaT = 0.15 and thetaB = 0.18; agreement ranged from 88.8% to 98.8%. Data correspond to Supplementary Fig. 1r.

*Excel file: Suppl_Table6_threshold_sensitivity.xlsx*

**Supplementary Table 7 | OPLS-DA coefficients and VIP scores for metabolomics features.** Worksheets: Bulk NEG, bulk brain negative-ion mode (112 features); Bulk POS, bulk brain positive-ion mode (45 features); Spatial SM, spatial cyst-neighbourhood negative-ion mode (183 features). Columns: m/z, OPLS-DA coefficient (CoeffCS), VIP, Compound ID (HMDB/LIPID MAPS, where available) and Description. The VIP = 1 threshold was used to define differential metabolites (Supplementary Table 4). Data correspond to Fig. 5c–h.

*Excel file: Suppl_Table7_VIP_analysis.xlsx*

## Supplementary Note 1 Annotation of host cell subclusters

### 1.1 Neuronal subclusters (Supplementary Fig. 1b)

The 27 neuronal subclusters comprise cerebellar granule cells (C-GC), highly plastic spiny neurons (HPSN), olfactory bulb granule cells (OB-GC), tectal sensory glutamatergic projection neurons (TS-GPN), perirhinal glutamatergic projection neurons (PRh-GPN), association cortical pyramidal neurons of layers II–IV (AC-GPN-2/4), direct-pathway D1 receptor-positive striatal medium spiny neurons (D1-MSN), highly active spiny neurons (HASN), association cortical pyramidal neurons of layers II–III (AC-GPN-2/3), indirect-pathway D2 receptor-positive striatal medium spiny neurons (D2-MSN), dentate gyrus granule cells (DG-GC), association cortical pyramidal neurons of layers V–VI (AC-GPN-5/6), hippocampal CA1/CA3 pyramidal neurons (Hipp-GPN-CA1/3), SST^+^ interneurons (SST^+^ IN), basket cells (BC), neuroblasts (NB), periglomerular cells (PGC), VIP^+^ GABAergic interneurons (VIP^+^ IN), hybrid-phenotype young dopaminergic neurons (HP-yDA), short-axon cells (SAC), chandelier cells (ChC), nucleus accumbens-derived direct-pathway D1 medium spiny neurons with Drd3 co-expression (Drd3 + NAc_D1-MSN), Purkinje cells (PkC), mitral cells (MC), circuit-integrating neurons (CIN), thalamic glutamatergic projection neurons (Thal-GPN) and glycinergic interneurons (GlyIN). Glutamatergic neurons such as cortical pyramidal neurons (AC-GPNs-2/3, AC-GPNs-2/4, AC-GPNs-5/6) and hippocampal pyramidal neurons (Hipp-GPNs-CA1/3) decreased significantly in proportion, as did GABAergic neurons such as striatal D1-MSNs, D2-MSNs and VIP^+^ INs, and olfaction-related cells (OB-GCs, PRh-GPNs, PGCs, SACs, MCs). The marked decline of neuroblasts (NBs) and immature neurons (HP-yDA) suggests that *T. gondii* infection may impair adult neurogenesis. The two functionally defined neuronal subclusters identified by unsupervised clustering, highly active spiny neurons (HASN) and highly plastic spiny neurons (HPSN), showed significant increase and decrease, respectively, indicating reduced neuronal plasticity upon infection; the increased proportion of HASNs during chronic infection may relate to parasite-promoted neuronal dopamine production. The increase in circuit-integrating neurons (CINs) may suggest that infection promotes neuronal circuit re-integration.

### 1.2 Astrocyte subclusters (Supplementary Fig. 1c)

The eight astrocyte subclusters comprise homeostatic astrocytes 1–3 (HA1, HA2, HA3), interferon responsive astrocytes (IRA), transitional reactive astrocytes (TRA), immune effector astrocytes (IEA), vascular-associated astrocytes (VA) and oxidative stress-responsive astrocytes (ORA).

### 1.3 Microglial subclusters (Supplementary Fig. 1d)

The five microglial subclusters comprise interferon-responsive microglia (IRM), homeostatic microglia 1 and 2 (HM1, HM2), proliferative microglia (PM) and pro-inflammatory microglia (PIM). As brain-resident immune cells, microglia respond rapidly to brain injury or pathogen infection. Microglial numbers increased significantly during acute *T. gondii* infection and remained elevated through the chronic phase. Homeostatic microglia (HM1 and HM2) decreased significantly after infection, whereas IRM, PM and PIM increased; homeostatic microglial numbers partially recovered during chronic infection. Pseudotime analysis showed that early-pseudotime cells were mainly homeostatic microglia, predominantly from uninfected mice; as infection proceeded, homeostatic microglia differentiated into IRM, PM and PIM pro-inflammatory/proliferative states.

### 1.4 Monocyte/macrophage subclusters (Supplementary Fig. 1e)

The five monocyte/macrophage subclusters comprise classical monocytes (cMo), Tgfbr1^+^ monocyte-derived macrophages (Tgfbr1^+^ MoDM), M1 monocyte-derived macrophages (Mo-M1 Mac), regulatory monocyte-derived macrophages (Mo-Mreg) and border-associated macrophages (BAM). Among peripheral immune cells, monocytes/macrophages are rapidly recruited to the CNS and infiltrate the brain parenchyma after *T. gondii* infection; brain-resident border-associated macrophages also contribute to pathogen defense. Driven by peripheral and BBB-associated pro-inflammatory cues and by direct parasite stimuli, circulating monocytes progressively differentiate into macrophages that control *T. gondii* infection through antigen presentation and pro-/anti-inflammatory functions.

### 1.5 Dendritic cell subclusters (Supplementary Fig. 1f)

The four dendritic cell subclusters comprise conventional dendritic cells type 1 (cDC1) and type 2 (cDC2), migratory dendritic cells (migDC) and plasmacytoid dendritic cells (pDC).

### 1.6 T cell subclusters (Supplementary Fig. 1g)

The nine T cell subclusters comprise CD8^+^ T cells 1 (CD8^+^ T 1), CD4^+^ T cells/T follicular helper cells(CD4^+^ T/Tfh), activated CD8^+^ cytotoxic T cells/natural killer T cells (Act CD8^+^ T/NKT), exhausted/proliferating CD8^+^ T cells (Tex/Tpro), γδT cells (γ δT), CD8^+^ T cells 2 (CD8^+^ T 2), regulatory T cells (Treg), T helper 2 cells (TH2) and IL7R^+^ memory T cells (IL7R^+^ Tm). Among peripheral immune cells, T cells are rapidly recruited to the CNS and infiltrate the brain parenchyma after *T. gondii* infection. T cells are extremely rare in uninfected mouse brain, increase markedly during acute and chronic infection, and differentiate into functionally distinct subclusters, further supporting their important role in anti-*T. gondii* immunity during acute and chronic infection.

### 1.7 B cell and plasma cell subclusters (Supplementary Fig. 1h, i)

The four B cell subclusters comprise naive B cells (Naive B), activated B cells (Act B), germinal center B cells (GC B) and memory B cells (Mem B). The two plasma cell subclusters comprise mature plasma cells (MPC) and plasmablasts (PB). Unlike T cells, B cells and plasma cells were not recruited to the brain during acute infection: they were identified only in chronically infected brains, and plasma cells were entirely absent from uninfected mice. Beyond producing antibodies against *T. gondii*, B cells have recently been shown to maintain the architecture of the antigen-presenting cell network, indirectly determining the efficiency of CD8^+^ T cell priming, which may partly explain their marked increase during chronic *T. gondii* infection.

### 1.8 Fibroblast subclusters (Supplementary Fig. 1j)

The five fibroblast subclusters comprise myofibroblasts (MFB), lymphocyte-interacting fibroblasts (LIFB), dural fibroblasts (DFB), inflammatory fibroblasts (IFB) and leptomeningeal fibroblasts (LFB).

### 1.9 Mural/endothelial cell subclusters (Supplementary Fig. 1k)

The three mural/endothelial subclusters comprise endothelial cells (EC), pericytes (Peri) and vascular smooth muscle cells (VSMC).

## Supplementary Methods 1 LC–MS mobile-phase gradient and mass-spectrometry parameters

Bulk metabolomics was performed on a Waters ACQUITY Premier ultra-high-performance liquid chromatography (UPLC) system coupled to a quadrupole time-of-flight mass spectrometer, equipped with an ACQUITY Premier BEH C18 1.7 μm VanGuard FIT column (2.1 × 100 mm). Phase A was 60% acetonitrile/40% water and phase B was 90% isopropanol/10% acetonitrile, both containing 10 mM ammonium formate and 0.1% formic acid. The mobile-phase gradient is given in Table 1 and the mass-spectrometry parameters in Table 2.

## References

1. Montoya, J. G. & Liesenfeld, O. Toxoplasmosis. Lancet 363, 1965–1976 (2004).

2. Pappas, G., Roussos, N. & Falagas, M. E. Toxoplasmosis snapshots: global status of *Toxoplasma gondii* seroprevalence and implications for pregnancy and congenital toxoplasmosis. Int. J. Parasitol. 39, 1385–1394 (2009).

3. Elsheikha, H. M., Marra, C. M. & Zhu, X.-Q. Epidemiology, pathophysiology, diagnosis, and management of cerebral toxoplasmosis. Clin. Microbiol. Rev. 34, e00115–19 (2020).

4. Hunter, C. A. & Sibley, L. D. Modulation of innate immunity by *Toxoplasma gondii* virulence effectors. Nat. Rev. Microbiol. 10, 766–778 (2012).

5. Ramírez-Flores, C. J. & Mondragón-Flores, R. Comprehensive analysis of *Toxoplasma gondii* migration routes and tissue dissemination in the host. PLOS Negl. Trop. Dis. 19, e0013369 (2025).

6. Ferguson, D. J. P. & Hutchison, W. M. The host-parasite relationship of *Toxoplasma gondii* in the brains of chronically infected mice. Virchows Arch. A Pathol. Anat. Histopathol. 411, 39–43 (1987).

7. Blanchard, N., Dunay, I. R. & Schlüter, D. Persistence of *Toxoplasma gondii* in the central nervous system: a fine-tuned balance between the parasite, the brain and the immune system. Parasite Immunol. 37, 150–158 (2015).

8. Tyebji, S., Seizova, S., Hannan, A. J. & Tonkin, C. J. Toxoplasmosis: a pathway to neuropsychiatric disorders. Neurosci. Biobehav. Rev. 96, 72–92 (2019).

9. Torrey, E. F., Bartko, J. J. & Yolken, R. H. *Toxoplasma gondii* and other risk factors for schizophrenia: an update. Schizophr. Bull. 38, 642–647 (2012).

10. Burgdorf, K. S. et al. Large-scale study of *Toxoplasma* and Cytomegalovirus shows an association between infection and serious psychiatric disorders. Brain Behav. Immun. 79, 152–158 (2019).

11. Nayeri Chegeni, T., et al. Is *Toxoplasma gondii* a potential risk factor for Alzheimer’s disease? A systematic review and meta-analysis. Microb. Pathog. 137, 103751 (2019).

12. Kavakebian, M. et al. The cognitive and neurological implications of *Toxoplasma gondii* infection: evidence from a systematic review and meta-analysis. Acta Trop. 268, 107705 (2025).

13. Miman, O., Kusbeci, O. Y., Aktepe, O. C. & Cetinkaya, Z. The probable relation between *Toxoplasma gondii* and Parkinson’s disease. Neurosci. Lett. 475, 129–131 (2010).

14. Kamal, A. M., Kamal, A. M., Abd El-Fatah, A. S., Rizk, M. M. & Hassan, E. E. Latent toxoplasmosis is associated with depression and suicidal behaviour. Arch. Suicide Res. 26, 819–830 (2022).

15. Markovitz, A. A., et al. *Toxoplasma gondii* and anxiety disorders in a community-based sample. Brain Behav. Immun. 43, 192–197 (2015).

16. De Haan, L., Sutterland, A. L., Schotborgh, J. V., Schirmbeck, F. & De Haan, L. Association of *Toxoplasma gondii* seropositivity with cognitive function in healthy people: a systematic review and meta-analysis. JAMA Psychiatry 78, 1103–1112 (2021).

17. Navia, B. A. et al. Cerebral toxoplasmosis complicating the acquired immune deficiency syndrome: clinical and neuropathological findings in 27 patients. Ann. Neurol. 19, 224–238 (1986).

18. Porter, S. B. & Sande, M. A. Toxoplasmosis of the central nervous system in the acquired immunodeficiency syndrome. N. Engl. J. Med. 327, 1643–1648 (1992).

19. Robert-Gangneux, F. & Dardé, M.-L. Epidemiology of and diagnostic strategies for toxoplasmosis. Clin. Microbiol. Rev. 25, 264–296 (2012).

20. Kanyama, C. et al. Causes of HIV-related CNS infection in Cameroon, Malawi, and Tanzania: epidemiological findings from the DREAMM HIV-related CNS implementation study. Lancet Glob. Health 13, e345–e354 (2025).

21. Vastava, P. B. et al. MRI features of *Toxoplasma* encephalitis in the immunocompetent host: a report of two cases. Neuroradiology 44, 834–838 (2002).

22. Giannakis, A., Makridimas, M., Christaki, E., Zikou, A. K. & Konitsiotis, S. Central nervous system toxoplasmosis with atypical neurologic findings in an immunocompetent patient. Lab. Med. 57, lmaf068 (2026).

23. Beltrame, A., Kim, K. & Weiss, L. M. Human toxoplasmosis: current advances in the field. Curr. Opin. Infect. Dis. 38, 392–402 (2025).

24. Rovito, D., Zanet, S., Trunfio, M. G. C., Calcagno, A. & Ferroglio, E. Neurocognitive effects of latent *Toxoplasma gondii* infection in immunocompromised people with HIV. Parasitology advance online publication, 10.1017/S0031182025101509 (2026).

25. Alday, H. & Doggett, J. Drugs in development for toxoplasmosis: advances, challenges, and current status. Drug Des. Devel. Ther. 11, 273–293 (2017).

26. Dubey, J. P., Lindsay, D. S. & Speer, C. A. Structures of *Toxoplasma gondii* tachyzoites, bradyzoites, and sporozoites and biology and development of tissue cysts. Clin. Microbiol. Rev. 11, 267–299 (1998).

27. Cabral, C. M. et al. Neurons are the primary target cell for the brain-tropic intracellular parasite *Toxoplasma gondii*. PLOS Pathog. 12, e1005447 (2016).

28. Dzierszinski, F., Nishi, M., Ouko, L. & Roos, D. S. Dynamics of *Toxoplasma gondii* differentiation. Eukaryot. Cell 3, 992–1003 (2004).

29. Ferguson, D. J. P., Hutchison, W. M. & Pettersen, E. Tissue cyst rupture in mice chronically infected with *Toxoplasma gondii*: an immunocytochemical and ultrastructural study. Parasitol. Res. 75, 599–603 (1989).

30. Hermes, G. et al. Neurological and behavioural abnormalities, ventricular dilatation, altered cellular functions, inflammation, and neuronal injury in brains of mice due to common, persistent, parasitic infection. J. Neuroinflammation 5, 48 (2008).

31. Schaeffer, M. et al. Dynamic imaging of T cell-parasite interactions in the brains of mice chronically infected with *Toxoplasma gondii*. J. Immunol. 182, 6379–6393 (2009).

32. Zhao, X.-Y. & Ewald, S. E. The molecular biology and immune control of chronic *Toxoplasma gondii* infection. J. Clin. Invest. 130, 3370–3380 (2020).

33. Koshy, A. A., et al. *Toxoplasma* secreting Cre recombinase for analysis of host-parasite interactions. Nat. Methods 7, 307–309 (2010).

34. Yamashita, K., Yui, K., Ueda, M. & Yano, A. Cytotoxic T-lymphocyte-mediated lysis of *Toxoplasma gondii*-infected target cells does not lead to death of intracellular parasites. Infect. Immun. 66, 4651–4655 (1998).

35. Suzuki, Y. et al. Removal of *Toxoplasma gondii* cysts from the brain by perforin-mediated activity of CD8^+^ T cells. Am. J. Pathol. 176, 1607–1613 (2010).

36. Ferguson, D. J. P., Graham, D. I. & Hutchison, W. M. Pathological changes in the brains of mice infected with *Toxoplasma gondii*: a histological, immunocytochemical and ultrastructural study. Int. J. Exp. Pathol. 72, 463– 474 (1991).

37. Dotiwala, F. et al. Killer lymphocytes use granulysin, perforin and granzymes to kill intracellular parasites. Nat. Med. 22, 210–216 (2016).

38. Prandovszky, E. et al. The neurotropic parasite *Toxoplasma gondii* increases dopamine metabolism. PLOS ONE 6, e23866 (2011).

39. Calvo-Urbano, B. et al. The role of parasite-produced dopamine in *Toxoplasma gondii*-altered host behaviour. Nat. Commun. 16, 10906 (2025).

40. Wang, Y. et al. Innate immune and metabolic signals induce mitochondria-dependent membrane lysis via mitoxyperiosis. Cell 188, 7155–7174.e25 (2025).

41. Salvioni, A. et al. Robust control of a brain-persisting parasite through MHC I presentation by infected neurons. Cell Rep. 27, 3254–3268.e8 (2019).

42. Porte, R., et al. Protective function and differentiation cues of brain-resident CD8^+^ T cells during surveillance of latent *Toxoplasma gondii* infection. Proc. Natl. Acad. Sci. U.S.A. 121, e2403054121 (2024).

43. Koshy, A. A., et al. *Toxoplasma* co-opts host cells it does not invade. PLOS Pathog. 8, e1002825 (2012).

44. Tabaie, E. Z., et al. *Toxoplasma gondii* infection of neurons alters the production and content of extracellular vesicles directing astrocyte phenotype and contributing to the loss of GLT-1 in the infected brain. PLOS Pathog. 21, e1012733 (2025).

45. Latifi, A. & Flegr, J. Beyond latency: chronic *Toxoplasma* infection and its unveiled behavioural and clinical manifestations—a 30-year research perspective. Biomedicines 13, 1731 (2025).

46. Melzer, T. C., Cranston, H. J., Weiss, L. M. & Halonen, S. K. Host cell preference of *Toxoplasma gondii* cysts in murine brain: a confocal study. J. Neuroparasitology 1, 1–6 (2010).

47. Figueiredo, C. A. et al. Immune response and pathogen invasion at the choroid plexus in the onset of cerebral toxoplasmosis. J. Neuroinflammation 19, 17 (2022).

48. Schneider, C. A., et al. Imaging the dynamic recruitment of monocytes to the blood–brain barrier and specific brain regions during *Toxoplasma gondii* infection. Proc. Natl. Acad. Sci. U.S.A. 116, 24796–24807 (2019).

49. Olivera, G. C., Ross, E. C., Peuckert, C. & Barragan, A. Blood-brain barrier-restricted translocation of *Toxoplasma gondii* from cortical capillaries. eLife 10, e69182 (2021).

50. Rodriguez, M. E. et al. ICAM-1/CD18-mediated sequestration of parasitised phagocytes in cortical capillaries promotes neuronal colonisation by *Toxoplasma gondii*. Nat. Commun. 16, 3529 (2025).

51. De Castro, L. L., et al. *Toxoplasma gondii* impairs CX3CL1/fractalkine shedding from mouse cortical neurons, leading to microglia activation. Microbiol. Spectr. 13, e01074–25 (2025).

52. Parlog, A. et al. Chronic murine toxoplasmosis is defined by subtle changes in the neuronal connectivity. Dis. Model. Mech. 7, 459–469 (2014).

53. Ihara, F., et al. *Toxoplasma gondii* infection in mice impairs long-term fear memory consolidation through dysfunction of the cortex and amygdala. Infect. Immun. 84, 2861–2870 (2016).

54. Tao, Q., et al. *Toxoplasma gondii* Chinese I genotype Wh6 strain infection induces tau phosphorylation via activating GSK3β and causes hippocampal neuron apoptosis. Acta Trop. 210, 105560 (2020).

55. Torres, L., et al. *Toxoplasma gondii* alters NMDAR signalling and induces signs of Alzheimer’s disease in wild-type, C57BL/6 mice. J. Neuroinflammation 15, 57 (2018).

56. Mahmoud, M. E., Fereig, R. & Nishikawa, Y. Involvement of host defense mechanisms against *Toxoplasma gondii* infection in anhedonic and despair-like behaviours in mice. Infect. Immun. 85, e00007–17 (2017).

57. Mahmoud, M. E., Ihara, F., Fereig, R. M., Nishimura, M. & Nishikawa, Y. Induction of depression-related behaviours by reactivation of chronic *Toxoplasma gondii* infection in mice. Behav. Brain Res. 298, 125–133 (2016).

58. Sibley, L. A. et al. Caspase-8 expression in CD8^+^ T cells promotes pathogen restriction in the brain during *Toxoplasma gondii* infection. Sci. Adv. 11, eadz4468 (2025).

59. Belloy, M., et al. *Toxoplasma gondii* infection and chronic IL-1 elevation drive hippocampal DNA double-strand break signalling, leading to cognitive deficits. Nat. Neurosci. 28, 2067–2077 (2025).

60. Suzuki, Y., Claflin, J., Wang, X., Lengi, A. & Kikuchi, T. Microglia and macrophages as innate producers of interferon-gamma in the brain following infection with *Toxoplasma gondii*. Int. J. Parasitol. 35, 83–90 (2005).

61. Zhang, Y. et al. Activated microglia contribute to neuronal apoptosis in toxoplasmic encephalitis. Parasit. Vectors 7, 372 (2014).

62. David, C. N. et al. GLT-1-Dependent disruption of CNS glutamate homeostasis and neuronal function by the protozoan parasite *Toxoplasma gondii*. PLOS Pathog. 12, e1005643 (2016).

63. Ulu, A. et al. Bradyzoite subtypes rule the crossroads of *Toxoplasma* development. Nat. Commun. 17, 1783 (2026).

64. Giuliano, C. J. et al. Convergent evolution of metabolic regulation governs redox adaptation in *Toxoplasma*. Cell 189, 6338–6355.e10 (2026).

65. Wu, Q.-Q. et al. Uromodulin p.His36Tyr promotes macrophage pyroptosis via App-Cd74 signalling to drive renal inflammation in ADTKD. Nat. Commun. 17, 5863 (2026).

66. Zhang, W. et al. Spatial and single-cell transcriptomics reveals senescence-associated changes in MIA-induced ASD male mouse brain. Cell Rep. 45, 117191 (2026).

67. Dos Reis, R. S., et al. Microglial reactivity and nodule formation are associated with synaptodendritic damage in the brains of people with HIV-1. Brain Pathol. 36, e70064 (2026).

68. Böyum, A. Isolation of mononuclear cells and granulocytes from human blood. Isolation of monuclear cells by one centrifugation, and of granulocytes by combining centrifugation and sedimentation at 1 g. Scand. J. Clin. Lab. Invest. Suppl. 97, 77–89 (1968).

69. Cornelissen, A. W. C. A., Overdulve, J. P. & Hoenderboom, J. M. Separation of *Isospora (Toxoplasma) gondii* cysts and cystozoites from mouse brain tissue by continuous density-gradient centrifugation. Parasitology 83, 103– 108 (1981).

70. Liao, S. et al. Stereo-cell: spatial enhanced-resolution single-cell sequencing with high-density DNA nanoball-patterned arrays. Science 389, eadr0475 (2025).

